# Spruce hybrids show superior cumulative growth and intermediate response to climate anomalies, as compared to their ecologically divergent parental species

**DOI:** 10.1101/2024.12.19.629043

**Authors:** Edouard Reed-Métayer, Claire Depardieu, Patrick Lenz, Jean Bousquet, Martin Perron

## Abstract

Climate change brings new constraints to which trees will have to adapt, including more frequent weather extremes. Black spruce and red spruce are phylogenetically close but adapted to different ecological conditions, and they form a natural hybrid zone where their distributions come into contact. Thus, they represent an interesting model to study the effect of introgressive hybridization in the context of climate change, given that interspecific gene flow could affect their capacity to adapt where their natural distributions overlap. Using a common-garden field test gathering 20-year-old progeny trees resulting from rigorous controlled crosses including previously verified genetic identity of the parents, growth patterns and wood density differences could be observed between species and between them and their F_1_ hybrids. A dendroecological analytical approach relying on wood cores was used and revealed similar wood responses to climatic variations between species, both through lifetime climate sensitivity and through episodic stress response indexes. They were however differentially expressed in early- and latewood between black spruce and red spruce, differences likely driven by diverging cambial phenology adaptations to different growing season lengths. F_1_ hybrids exhibited hybrid vigor for cumulative growth under the test site conditions, but showed intermediate values for traits related to climate response. These results offer new perspectives for understanding the dynamics of adaptation in hybrid zones in the context of climate change, as well as for guiding conservation and genetic improvement efforts.

**Highlights:**

1. F_1_ cumulative growth is higher than their ecologically contrasted parents.
2. Hybrids had intermediate climate responses
3. Parental species responded similarly to climate anomalies, but in various intra-ring part.
4. Cambial phenology likely drives divergences in wood reaction to climate anomalies

## 1. Introduction

Globally, forests are increasingly facing abiotic and biotic stresses due to climate change, leading to reduced resilience and ultimately higher mortality rates (Seidl et al., 2017; Forzieri et al., 2022; Hartmann et al., 2022). As the Earth’ climate is on its way to reach a 1.5°C temperature increase by the early 2030s, it coincides with the likely emergence of irreversible tipping points (Dakos et al., 2019; IPCC, 2022). As a result, the increasing frequency and intensity of severe climatic episodes such as droughts, early and late frosts, heatwaves, and forest fires will impact even more tree adaptation, productivity, and survival (Knapp et al., 2002; Sass-Klaassen et al., 2016; Ammer, 2019). Both cold and drought episodes exert a major influence on tree productivity and physiology in temperate and boreal regions, albeit through different mechanisms (Harfouche et al., 2014; Vitasse et al., 2019). Cold temperatures, when mismatched with phenological rhythms due to temperature fluctuations, can lead to immediate impacts on growth (Dittmar et al., 2006; Benomar et al., 2022). As for droughts, they induce water deficits within the trees, causing hydraulic imbalance and hence a reduction in photosynthesis. This can, in turn, result in a sustained decrease in radial growth and, ultimately, in increasing tree mortality, especially when drought episodes occur repeatedly (Grote et al., 2016; Choat et al., 2018; Gentilesca et al., 2021).

Natural tree populations exhibit population-specific climate adaptation (Morgenstern, 1996), and they show varying levels of genetically-determined ecophysiological predispositions to phenotypic plasticity and response in facing climatic conditions (Benomar et al., 2015, 2016; Berlin et al., 2017). These population-specific climate adaptations, along with generally high levels of genetic diversity, used to maintain a dynamic equilibrium resulting in adequate health, survival, and productivity of tree populations under similar climates, including climate anomalies, thus seed source genetic quality and matching to site has had high importance in reforestation contexts both historically and currently (Morgenstern, 1996). But as the contemporary climate change takes place at an unprecedented rate and induces rapidly changing climatic conditions and more important stress episodes, natural and reforested tree populations can be taken short, if a lack of sufficient adaptive genetic variation or acclimation responses to new levels of climate anomalies occurs (Benomar et al., 2022). Assessing adaptative variation of genetic nature and acclimatization responses, considering local environmental pressures, and understanding the genetic factors behind climate adaptation are therefore essential for sustainable forest management and dynamic conservation efforts (Howe & Brunner, 2005; Savolainen et al., 2013; Otis-Prud’homme et al., 2018; Benomar et al. 2022).

As a special case of genetic variation occurring in some tree species-pairs, natural introgressive hybridization is an additional generator of genotypic diversity (Gérardi et al., 2010; Godbout et al., 2012). Although the subject of debates in the now rapidly evolving perspective of dynamic ecological conservation and forest management in the context of climate change, hybridization could be considered in some cases not as an enemy to pure lineages, but as an opportunity of improving the adaptive potential of involved populations facing changing environmental conditions, attributable to the increase in genetic diversity (Allendorf et al., 2001; Quilodrán et al., 2020). Indeed, hybridization and introgression have the potential to yield novel gene combinations and a mosaic of adaptive phenotypic traits from which natural selection or breeding can filter (Rieseberg et al., 1999; Gérardi et al., 2010; Godbout et al., 2012), making it a resource worth noting for the management and conservation of conifer genetic resources (Perron & Bousquet, 1997; de Lafontaine et al., 2015; Hamilton et al., 2016; De la Torre et al., 2017).

Limited studies on abiotic stress tolerance and climate adaptation traits in interspecific hybrids in tree species have however shown diverse inheritance patterns. Recent research found that naturally introgressed hybrids of two commercially significant pines (*P. contorta* × *P. banksiana*) displayed increased juvenile growth potential under normal and dry conditions, a result of the combination of the best parental characteristics (Bockstette et al., 2021). Conversely, introgressed hybrids of white (*Picea glauca*) and Engelmann spruce (*Picea engelmannii*) assessed in tree improvement programs showed higher survival in intermediate environments compared to parental species (De La Torre et al., 2014). In a study of hybrid taxa of *Pinus elliottii* (Engelman) and *Pinus caribaea* (Morelet), hybrids deviated away from the mid-parent value and toward the parent species more susceptible to freezing (Duncan et al., 1996). Hybrids of *Juglans cinerea* and *Juglans ailantifolia* exhibited intermediate cold and heat tolerance compared to their progenitors (Brennan et al., 2021). Given these highly variable observations, further research is needed to understand the role of hybridization in shaping climate-related traits in trees, and to better understand vulnerability and resilience to climate shifts and instability in species complexes.

Dendroecology appears as a well-suited investigative tool to better understand tree responses to climatic stress periods, as it allows the retrospective analysis of their effects on tree productivity, physiology and other related processes. Indeed, under the conditions where climate induces a stress, a limitation, or any measurable biological response to the trees, the analysis of wood increment cores can indeed reveal past acclimation strategies (Housset et al., 2018; D’Andrea et al., 2020; Zheng et al., 2022). Recent studies have furthermore shown that the use of tree-ring traits reflecting tree responses to climatic constraints in experimental setups designed for the analysis of genetic effects could enhance our understanding of the genetic basis and underlying physiological processes related to cold or drought responses (Heer et al., 2018; Housset et al., 2018; Depardieu et al., 2021). Notably, climate-sensitivity traits, which assess how a wood trait responds to a single climate variable throughout a tree’s lifespan, have allowed the discovery of genes related to cold adaptation in *Pinus strobus* (Housset et al., 2018) and drought adaptation in *Picea glauca* (Depardieu et al., 2021). The plasticity of wood anatomical characteristics in response to both occasional and severe climatic stress periods, encompassing resistance, recovery, and overall resilience (Lloret et al., 2011), has also deepened our understanding of the physiological mechanisms involved in reacting to cold and drought (Isaac-Renton et al., 2018; Matisons et al., 2019; Depardieu et al., 2020). Significant heritability estimates of resilience to drought have also been reported, thus suggesting a partially inherited capacity for plastic acclimatization responses to drought throughout the tree’s life (Depardieu et al., 2020; Laverdière et al., 2022). Dendroecological analysis therefore appears as a relevant and novel way to investigate genetic aspects of responses to climatic stress.

Black spruce (*Picea mariana* (Mill) B.S.P.) and red spruce (*Picea rubens* Sarg.) are phylogenetically closely related conifer species (Bouillé et al., 2011; Lockwood et al., 2013) and hybridize spontaneously in their zone of contact; yet they are ecologically very different species (Perron & Bousquet, 1997). Molecular studies suggest that red spruce likely originated from an ancestral black spruce population, isolated along the northeastern United States coast during a Pleistocene glaciation (Perron et al., 2000; de Lafontaine et al., 2015). Red spruce has a more limited natural distribution and exhibits lower genetic diversity than black spruce, likely due to a bottleneck effect during speciation (Perron et al., 2000), making it potentially more vulnerable in the context of adaptation to climate change (Prakash et al., 2022). While the individual evaluation of both species suggests they have low to moderate drought tolerance and are restricted by warm summers (Drobyshev et al., 2013; Ministère des Ressources Naturelles, 2013; Kosiba et al., 2018), it remains uncertain whether one species is more drought-tolerant than the other. Cold tolerance studies on the other hand show clearly marked differences between the two species, red spruce having lower cold hardiness compared to other ecologically similar mixed-forest conifers (DeHayes et al., 2001), and black spruce displaying a very high tolerance to cold temperatures (Strimbeck et al., 2015).

Despite the importance of understanding the adaptive response of both red spruce and black spruce and the impact of their introgressive hybridization in a changing climate, no research has yet rigorously compared those taxa’s responses to drought and cold periods using growth and wood traits as proxies for assessing their level of adaptation. Through a dendroecological approach, our study aimed to quantitatively compare growth, sensitivity traits to climate variation, and resilience to climatic stress periods in both red and black spruces, as well as in their hybrids. To achieve this, we employed sets of parental trees from various provenances to conduct both intra- and interspecific crosses, with species identity having been rigorously confirmed using species-specific molecular markers. F_1_ hybrids and pure species were then planted and grown during 20 years in a common garden study. We hypothesized that hybrid spruces would demonstrate intermediate growth patterns, sensitivity to climate variation and resilience compared to pure species, which implies higher growth, higher climate resiliency and lower climate vulnerability when compared to the least performing pure species for those traits. We also evaluated the climate impacts on both earlywood and latewood tree-ring series to provide a detailed characterization of the temporal influence of climate on wood growth and wood density. We postulated the presence of varying climate vulnerabilities among both wood series types and among the four groups (black spruce, red spruce, and their reciprocal F_1_ hybrid crosses) under examination.

## 2. Material and Methods

### 2.1. Biological material and progeny test

In spring 1994, the senior authors conducted controlled crosses between the monoecious black spruce and red spruce, as well as interspecific crosses in a reciprocal polycross mating design. Black spruce parental trees had their geographical origins mostly in the species allopatric area, thus largely avoiding the known zone of contact at the time. Red spruce parents, on the other hand, originated mostly from the northern part of its range but south of the central zone of contact. This range of geographic origins was the result of avoiding obviously unadapted red spruce material originating from too warm climates in the southern Appalachian Mountains of the United States, for the sake of ensuring a sufficient level of survival in the common garden located in the zone of contact, and ultimately producing data. Additionally, before conducting crosses, species identity of each black spruce or red spruce parent was validated using molecular markers developed by Perron et al. (1995) so to exclude any hybrid or introgressed parent tree of the mating design, making each polycross family derived from controlled crosses either pure red spruce, pure black spruce or F_1_ interspecific hybrid in either crossing direction (the two hybrid types were obtained). Each produced family was a cross between one mother parent and a custom polymix of 10 or 11 male (pollen) parents (also acting as mother parents in other families permitted by the monoecious nature of spruce species). For intraspecific crosses, a specific polymix was assembled to fertilize each mother tree so to exclude its own pollen and self-pollination. Spruce trees from 5 to 6 families for each of the four groups created (i.e. pure black spruce: *Pma*; pure red spruce: *Pru*; and both hybrid types: *Pma* × *Pru* and *Pru* × *Pma*) were included in the progeny test, for a total of 22 families (see **Table S1** for details). Seedlings spent their first 3 growing seasons under various light treatments in a greenhouse, after which they were outplanted in the field in the 1998 spring.

The progeny test was established in 1998 at Saint-Luc-de-Bellechasse, in southern Quebec, Canada (lat. 46°52ˈ N, long. 70°42ˈ W and alt. 480 m), in a region within the zone of contact (**Fig. 1**). The progeny test was planted on a recently harvested woodlot, with 3-tree row plots in a 24 randomized complete block design (RCBD). Blocks were organized perpendicularly to the light slope on which the progeny test was established, so to control for the main source of environmental site variation. Usual establishment and maintenance procedures were followed, including the replacement of dead trees in the first years following establishment of the field test, and use of planted border trees around the main blocks to keep light conditions as uniform as possible within the experimental blocks.

**Figure 1.**
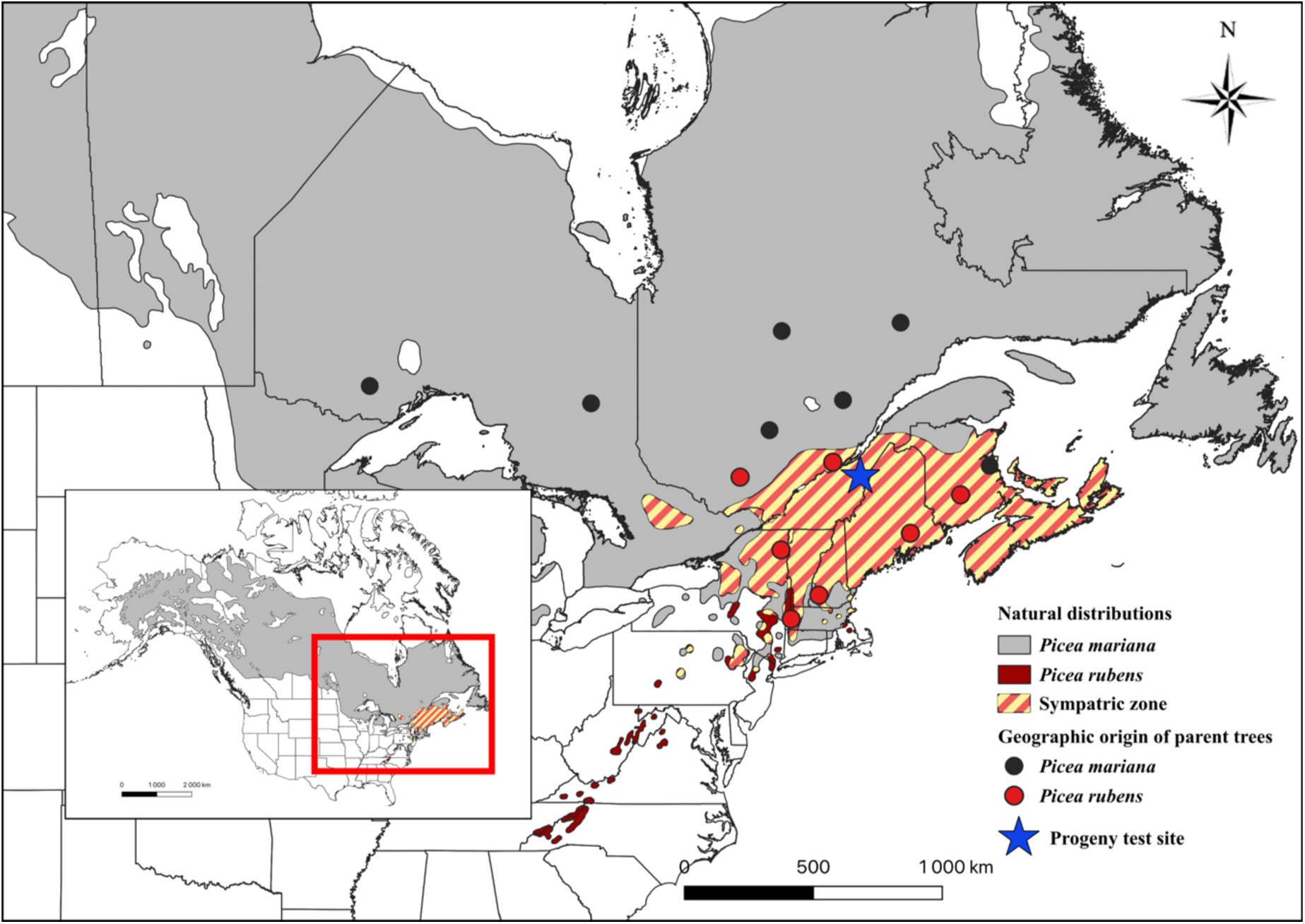
Progeny trial site, geographic origin of parent trees and the native ranges of red and black spruce. The progeny trial is marked by a blue star on the map and is located in southern Quebec, in the zone sympatric to the two species as depicted by the hatched pattern. The allopatric native distributions of black spruce and red spruce are respectively depicted in grey and dark red (data sources: Little, 1971; CEC, 2022).

### 2.2. Wood data collection and calculation of wood-related traits

Dendrometric traits such as height (H) and diameter at breast height (DBH, 1.3 m above ground) measurements were taken in autumn of 2007 and 2017, that is, 10 and 20 years after planting in the field. Wood increment cores were collected at breast height from the south side of trees in autumn of 2020. Wood cores were then planed to a thickness of approximately 1.7 mm and were kept in an 8% humidity conditioning room for two weeks prior to X-ray analysis. X-ray scanning of the dried increment cores was done with a QTRS-01X scanner of QMS (Quintek Measurement Systems Inc., Knoxville TN, USA) with a linear resolution of 0.02 mm, using the associated QMS tree ring system software.

Data extracted from the scanning tree core process included total ring, earlywood (ew), and latewood (lw) ring widths, as well as detailed wood density profiles for each of those tissues. Cross-dating of total ring width chronologies was carried out using COFECHA software (Holmes, 1983). Rings showing abnormal wood density profiles reflecting irregularities (such as resin pockets, small knots, etc.) and/or incompleteness due to the harvesting of cores during current-year growth were discarded for future analysis. Distinguishing between earlywood and latewood was done according to the mid-range method, where mid-range wood density of each ring (Dmr) is calculated as the mean between maximum (Dmax) and minimum wood density (Dmin) within each ring: Dmr = (Dmax+Dmin)/2 (Polge, 1978; Degron & Nepveu 1996). Transition from earlywood to latewood was then determined to be at the point where the last wood density yearly measurement lower than Dmr was followed by two successive yearly measurements higher than Dmr.

Wood-related traits such as basal area increment (BAI) and latewood percentage (LwP) were derived from ring width and wood density data. Ring widths were converted to BAI using the BAI.in function from dplR R package to directly reflect annual wood production (Bunn, 2010). In total, 961 trees were considered in the analyses.

### 2.3. Climate data

Daily climate data (maximum, minimum and mean temperature, precipitation sum) were obtained for the period 2003-2019 using the BioSIM software v.11.4 (Régnière et al., 2017). Daily climate data was interpolated from the four closest weather stations, adjusted for elevation and location differentials with regional gradients, and obtained by using the network of Environment Canada weather stations (Environment Canada, 2013), as well as from the Ministère de l’Environnement et de la Lutte contre les Changements Climatiques du Québec, the Centre informatique de prévision des ravageurs en agriculture, and the Solution-Mesonet weather station networks (Lepage & Bourgeois, 2011). Daily and/or monthly values of climate variables were then derived from interpolated climate data using monthly/annual means or sums. Indices such as maximum temperature (Tmax), minimum temperature (Tmin), frost days (FD) were generated using the software. Three climatic variables for assessing drought conditions in soil (soil moisture index, SMI), soil organic layers (drought code, DC) and air (Standardized Precipitation-Evaporation Index, SPEI) were used to identify drought episodes and testing for climate heterogeneity signals (**Table S2**; Begueria et al., 2014). The soil moisture index (SMI) was estimated for each month using the quadratic + linear formulation procedure, which accounts for water loss through evapotranspiration (simplified Penman–Monteith potential evapotranspiration) and water gain from precipitation (Hogg et al., 2013). Parameter values for maximum and critical available soil water were set at 300 mm and 400 mm, as previously described in Depardieu et al. (2020). The Standardized Precipitation-Evaporation Index, or SPEI, is a multiscalar drought index that was calculated by the difference between daily precipitation and daily potential evapotranspiration (PET) (Vincente-Serrano et al., 2012). The drought code (DC), a metric for rating the average moisture content of deep, compact organic layers (Van Wagner, 1987), was converted into a deficit index (1-DC values) to compare the direction and magnitude of drought during the studied period. Three climatic variables were also used for assessing cold spells or frosts, namely the late frost risk (LFR), the number of thaw days (WTD), and the number of extreme frost days (EFD). LFR corresponds to the degree-days above 5°C accumulated at time of last spring frost below 0°C. WTD is defined as the number of days with a minimum temperature above 0°C between December 15^th^ and March 15^th^ of the current winter. Finally, EFD is the number of days with a minimum temperature below - 30°C, from November of the previous winter to March of the current spring.

### 2.4. Detrending of tree-ring series and climate-traits correlations

To control for tree age and increasing densification of the common-garden test plantation over time, detrended individual chronologies of earlywood and latewood BAI and average wood density, along with latewood percentage, were computed with a spline function from dplR R package (Bunn, 2010). Mean-value chronologies per taxon were then calculated by averaging the rows of detrended data using Tukey’s biweight robust mean (function chron in the package dplR). Proxy-climate relationships were performed using the dcc function in the R package treeclim (Zang & Biondi, 2015). Climate-traits correlations were based on seasonal or annual values of climatic variables and annual values of wood traits for the 2003-2019 period. The seasonal values of the tested climatic variables were obtained by averaging the mean monthly values for spring (March, April, May), summer (June, July, August), autumn (September, October, November), and winter (December, January, February). A total of eighteen candidate climate variables were considered in this study for initial selection steps (see **Table S2** for all); only those that ended up most relevant for a particular type of climatic stress were kept for further analysis (see **Table 1**).

**Table 1.** Description and abbreviations of the climatic variables investigated in this study.

| Variable | Abbreviation (unit) | Definition |
| --- | --- | --- |
| Late frost risk | LFR | Degree days above 5°C accumulated at time of last spring frost below 0°C. |
| Number of thaw days | WTD | Number of days where minimum temperature is above 0°C, between December 15th and March 15th of the previous winter. |
| Extreme frost days | EFD | Number of days with minimum temperature below -30°C, from november to march of the previous winter. |
| Minimum value of daily minimum temperature | Tmin_season (°C) | Minimum temperature for a given season, as defined in the Climdex database (season abbreviations: Spr: spring; Sum: summer; Aut: autumn; Win: winter). Let $TN_n$ be the daily minimum temperatures in month $k$, period $j$. The minimum daily minimum temperature each month is then $TN_{nkj} = \min(TN_{nkj})$. |
| Maximum value of daily maximum temperature | Tmax_season (°C) | Maximum temperature for a given season, as defined in the Climdex database. Let $TX_x$ be the daily maximum temperatures in month $k$, period $j$. The maximum daily maximum temperature each month is then $TX_{xkj} = \max(TX_{xkj})$. |
| Standardized Precipitation Evapotranspiration Index | SPEI_season | Mean seasonal (autumn, spring and summer) standardized precipitation evapotranspiration index calculated according to the Tornwaithe method. |
| Soil moisture index | SMI_season (%) | Average soil moisture index over a given season (autumn, spring, or summer). |
| Summer cumulative degree-days | DD5 (°C) | Summer cumulative degree-days above 5°C. |

### 2.5. Estimation of climate-sensitivity traits

Climate-sensitivity (CS) traits were calculated as Pearson correlation coefficients between the tree-level detrended wood chronologies and each of the climate variables for which a significant signal of climate anomaly had been previously detected in the analyses of chronologies (see section 2.4 above). This category of tree-ring traits can theoretically be considered as a proxy of tree vulnerability to climate since it captures the overall effect of climatic constraints on annual fluctuations in wood parameters across the tree lifespan (Housset et al., 2018; Depardieu et al., 2021). More precisely, a CS trait value close to 1 or - 1 is indicative of a strong relationship between the wood trait and the climatic variable tested, while a value close to zero indicates no constraints imposed by the climatic variable on the trait tested. CS trait analysis required complete ring series, free of invalid rings caused by resin pockets or abnormalities, limiting the series to rings between two invalid ones. The 2009-2018 period was used to balance statistical power (621 individuals retained) and the analysis’s relevance by including as many years as possible.

### 2.6. Detection of pointer years and calculation of resilience components

To assess radial growth and wood density responses to climatic stress periods, a pointer-year analysis was performed on earlywood and latewood series of the traits studied, using the “pointRes” R package (van der Maaten-Theunissen et al., 2015; van der Maaten-Theunissen et al., 2021), along with a confirmation from visual inspection. In this study, we sought to quantify the amplitude of response of both growth (basal area increment) and wood traits (wood density, latewood percentage) to climatic stress periods detected in the analysis of chronologies. Components of tree resilience, namely resistance (Rs), recovery (Rc) and resilience (Rl), were calculated following the definition initially provided by Lloret et al. (2011) and extended to physiological-related tree-ring traits by Depardieu et al. (2024). The three components of resilience were computed based on the trend values of the characteristics tested for the early- and latewood series for BAI and WD, as well as for latewood percentage (**Fig. S1, Table S4**). Resistance (Rs) describes a tree’s ability to absorb changes in its wood or growth properties induced by a climatic stress, and was calculated as Dr/PreDr, where Dr and PreDr are the mean values of the trait considered during and before the stress period, respectively. Recovery (Rc) is defined as the tree’s ability to restore its initial wood properties or growth after stress, and was calculated as PostDr/Dr, where PostDr is the average value of the trait considered after the stress period. Resilience (Rl) reflects a tree’s ability to achieve growth rates or wood properties after the climatic stress period and was obtained by dividing the mean values of the post-stress trait by the mean values of the pre-stress trait. The resilience components (Rs, Rc, and Rl) were calculated for climatic stress periods that markedly impacted the studied wood traits, and for which phenotypic data were available to enable a precise assessment of both tree resistance and tree recovery after a stress period.

### 2.7. Statistical analyses

Linear mixed models were used to examine the effects of group, family, and time on the wood traits examined. Models were fitted using the ‘asreml’ function from the asreml-r v.4.0 R package (Butler et al., 2017) as follows:

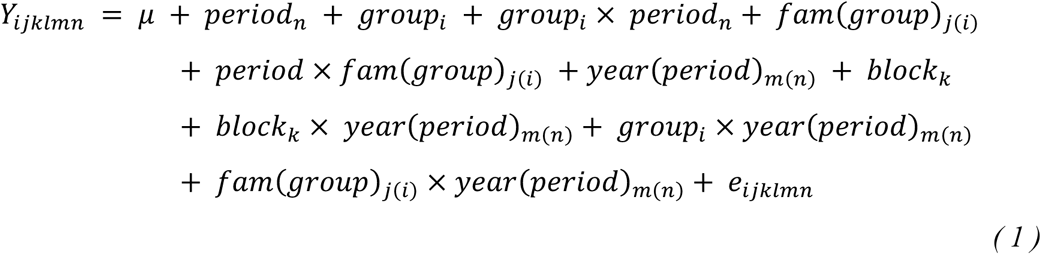

where *Y*_*ijklmn*_ is the tree-ring trait value on the *l*^th^ tree in the *m*^th^ year and *n*^th^ period from the *j*^th^ family within the *i*th group recorded in the *k*^th^ block; *µ* is the overall mean. *fam*(*group*)_*j*(*i*)_ is the family effect nested in *group*_*i*_, and *fam*(*group*)_*j*(*i*)_ and its interaction with the *period*_*n*_ effect were considered as fixed effects in the models. The timespan recorded in the tree-rings was divided in two distinct periods that represented initial growth (up to 2010) and a subsequent growth phase (after 2010) to easily evaluate the impact of time on the raw chronologies (**Fig. 2A-B**). The block effect and its interaction with the year (*year*_*m*_) were incorporated as random effects in the model. The effect of year nested within period (*year*(*period*)_*m*(*n*)_), the interaction of year and group (*group*_*i*_ × (*year*_*m*_)), as well as the interaction between year and family (*fam*(*group*)_*j*(*i*)_ × *year*_*m*_), were also included as random effects. *e* _*ijklmn*_ is the error term. Given the repeated measures on the same increment core for wood density and radial growth, the error term was assumed to be *e* ∼ *N* (0, *I*_*l*_ ⨂ *R*_*m*_), where *Il* is an identity matrix of dimension *l* (961 trees), *Rm* represents a first-order heterogeneous autoregressive correlation structure (AR1H) on time of dimension *m* × *m* (17 years × 17 years), and the symbol ⊗ refers to the Kronecker product. For the dendrometric traits, the CS traits, and the resilience traits that were not repeatedly measured over time, both the year effect and period and their specific interaction terms were omitted from the model, and the error term was specified as 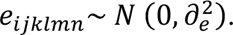 Data were transformed when normality and homogeneity of variances were not respected, and model parameters were estimated with the restricted maximum likelihood method (REML). A Tukey multiple comparison test was conducted, using the *posthoc* function from the biometryassist R package (Nielsen et al., 2023). Phenotypic correlations were computed as Pearson correlation coefficients, and their *p-*values were adjusted following Holm’s method using the *corr.test* function of the psych R package (Revelle, 2023). We compared dendrometric traits and basic wood variables (average BAI, wood density, etc.) with CS traits and resilience components with the goal of evaluating how growth trends and climate response traits related with growth and basic wood variables across and between each taxonomic group.

**Figure 2.**
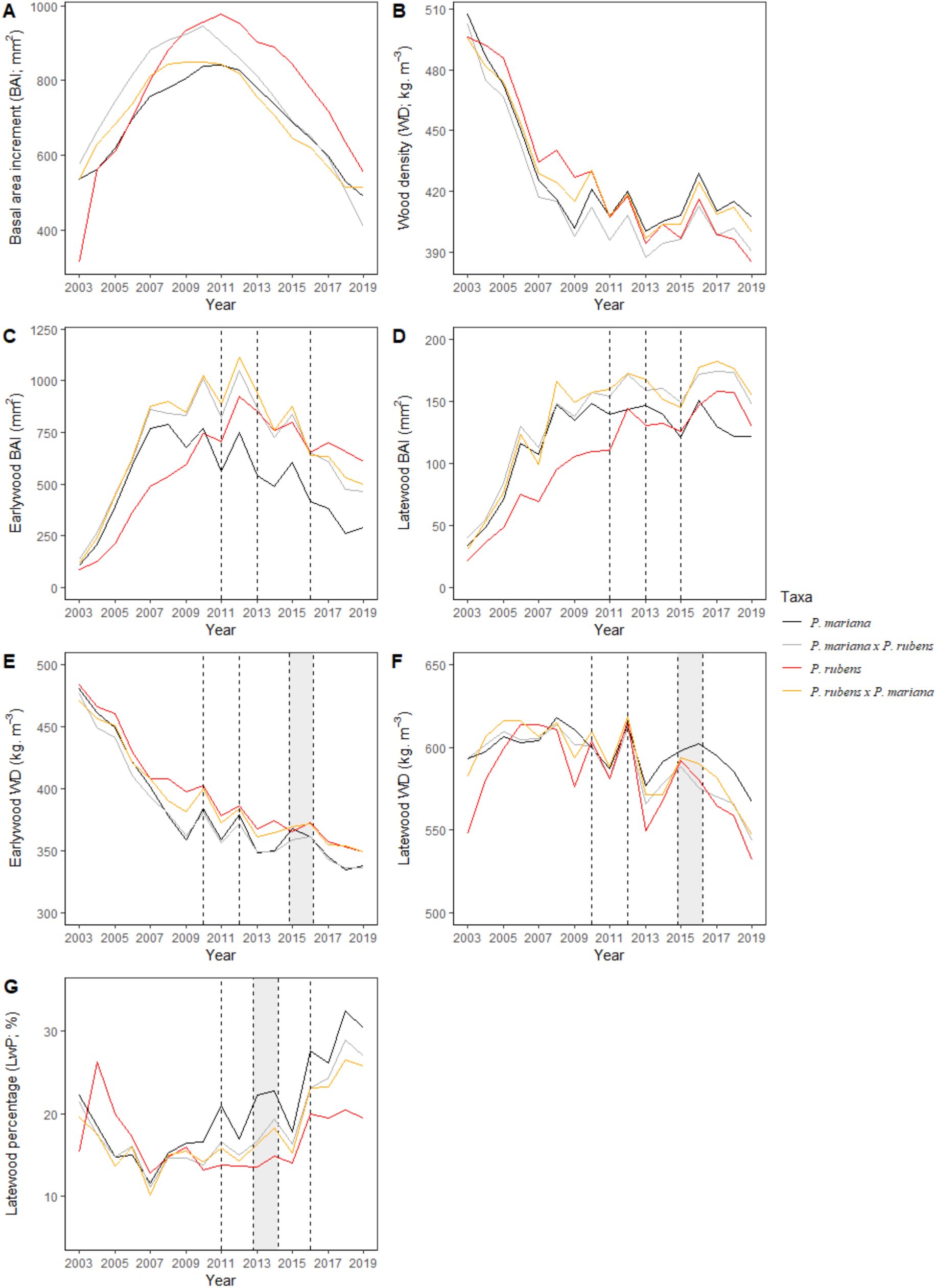
Inter-annual variation in radial growth, wood density and latewood percentage chronologies of black spruce, red spruce and their two reciprocal hybrid groups. Average time series by taxon are presented for the basal area increment (BAI; **A, C, E**) and the wood density (WD; **B, D, F**) at the whole ring (**A, B)**, the earlywood (**C, D**), and latewood levels (**E, F**). The latewood percentage (LwP) is presented in panel **G**. Both dashed lines and gray areas indicate selected years where wood parameters were considered altered by other factors than age- and growth-related trends (after detrending), for the 2003-2019 period. More precisely, the dashed lines indicate a specific year during which substantial variations were observed in the earlywood and latewood series as a result of drought and/or cold periods experienced by the trees. The gray areas indicate that multiple consecutive years were identified during which wood characteristics were significantly affected by the climate stress period under consideration.

## 3. Results

### 3.1. Lifetime trends of annual basal area increment and wood density

The time series of growth and wood density ring values for black spruce, red spruce and both hybrid types are shown in Figure 1. The lifetime trends of annual basal area increment and of wood density (**Fig. 2A-B**) reveal two distinct periods. The initial period spanning from 2003 to 2010 displays a collective rise in annual growth for all groups, accompanied by a gradual decrease in wood density across the groups. Subsequently, the second period from 2011 to 2019 demonstrated a gradual decline in annual growth rates, coupled with relatively stable wood density values. The four groups showed significant differences in the yearly trends for the growth for earlywood, latewood, and whole ring, with significantly different patterns over periods (**Table 2, Table S5**). Specifically, red spruce had slower earlier growth than black spruce and both hybrid types; however, it surpassed black spruce during the second period while both hybrid types remained the most rapidly growing (**Table 2**, **Fig. 2A-C-D**). This pattern is also reflected in cumulative growth, wherein both hybrid types and black spruce significantly exceeded red spruce in height and diameter until reaching at least 10 years in the field (**Table S6**). Wood density was significantly different among groups mostly during the second period, when red spruce and both hybrid types continued their decline to slightly lower wood density levels than black spruce (**Fig. 2B**, **Table 2**). When examined in terms of their earlywood and latewood components, the analysis revealed a significant effect of the group (ew *P* < 0.001, lw *P* = 0.0016), although the interaction between group and period was not significant (ew *P* = 0.12, lw *P* = 0.18) (**Table 2**).

**Table 2.**
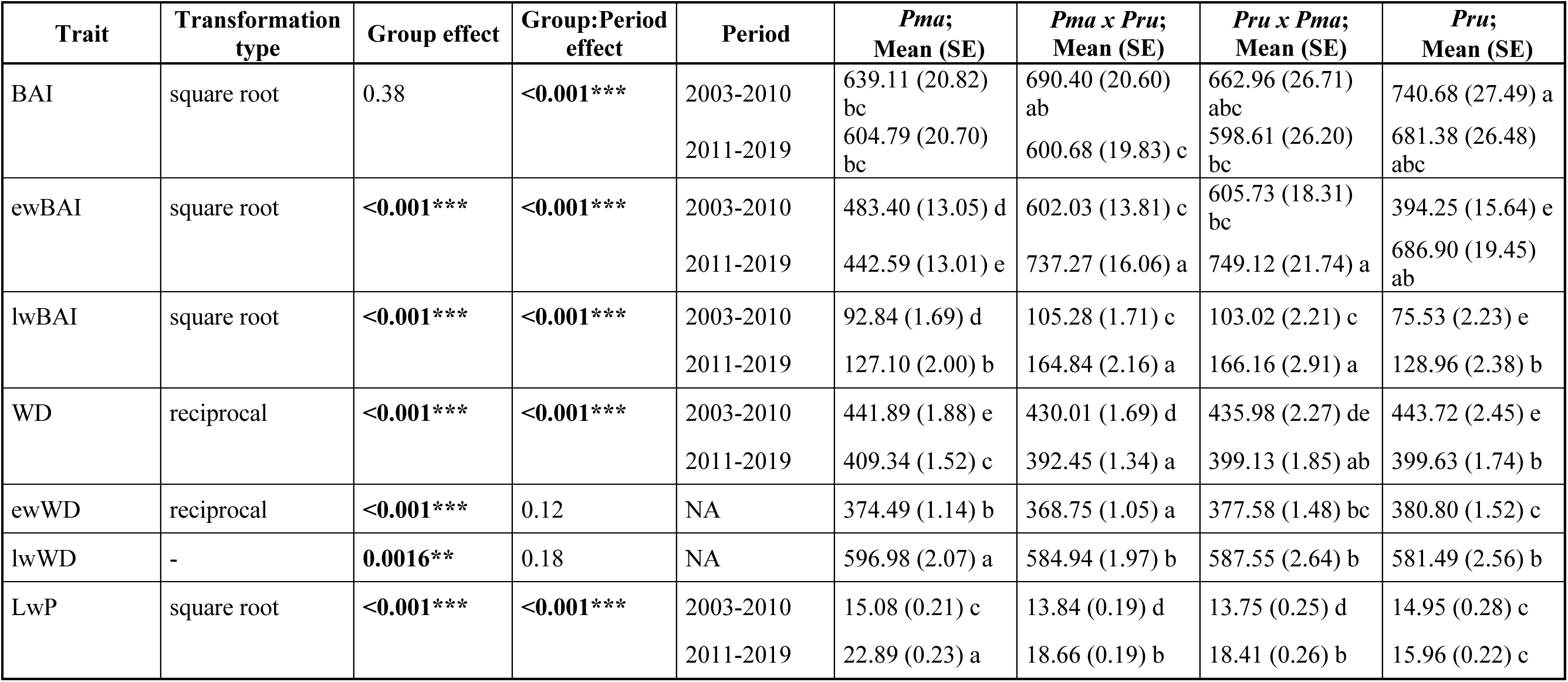
Linear modeling results for annual radial growth and wood density traits. The effects of group and the interaction between group and period effects were treated as fixed effects in the models. Period 1 refers to an early phase of tree growth between 2003 and 2010, while period 2 refers to the time period of tree growth from 2011 to 2019. Levels of significance for the effects tested are reported as follows:* *P* < 0.05, ** *P* < 0.01, and *** *P* < 0.001. Where the interaction effect (group:period) was significant (*P* < 0.05), post-hoc statistical grouping of groups was performed using Tukey’s pairwise comparison method with a significance level of 0.05, and significant differences between groups are indicated by different letters. BAI: basal area increment (whole-ring); ewBAI: earlywood basal area increment; lwBAI: latewood basal area increment; WD: wood density (whole-ring); ewWD: earlywood wood density; lwWD: latewood wood density; LwP: latewood percentage. In the case of the BAI, a non-significant group effect indicates that groups did not show significantly different growth over the entire study period; however, notable differences were observed between the two identified development phases or periods (i.e. 2002-2010 and 2011-2019).

Across groups, red spruce had the highest earlywood density, black spruce had the highest latewood density, and the hybrids showed intermediate or slightly lower wood density values compared to the pure species (**Table 2**, **Fig. 2E-F**). Over the course of time, latewood percentage exhibited an upward trend for all four groups (**Figure 2G**). While red spruce and black spruce displayed marginally latewood percentage than the hybrid types during the first period, black spruce exhibited a more pronounced increase compared to red spruce. In the second period, this ultimately led to higher latewood percentage for black spruce, with both hybrid types displaying intermediate values (**Fig. 2G**, **Table 2**).

### 3.2. Identification of drought and cold stress periods

The analysis of daily and seasonal temperatures revealed lower minimum temperatures during the winters of 2004, 2009, and 2014, as well as in the springs of 2015 and 2016 (**Fig. 3A-B)**. The period of cold stresses identified from 2014 to 2016 is therefore the combination of a particularly cold 2014 winter, cooler-than-average spring temperatures in 2015 and 2016, and higher count of thaw days during the 2016 winter (**Fig. 3B**). Otherwise, examining four drought indicators in conjunction with maximum temperatures has led us to discern three distinct drought periods that occurred either independently or in tandem with heat waves over the course of the study timespan (**Fig. 3A, 3C-D, Table S3**). A drought period occurred during the spring of 2010, evidenced by diminished soil moisture values and an increased number of dry days (**Fig. 3D**). A drought in 2012 exhibited pronounced soil drought conditions (**Fig. 3C**) and was further accentuated by a synchronous heat wave during the summer (**Fig. 3D**). The elevated count of dry days during this summer underscores the prolonged nature of the drought conditions (**Fig. 3D**). Furthermore, the 2012 drought conditions may have been enhanced by an exceptional episode of winter thaw in late March of the same year, most probably reducing water availability early in the growing season due to the reduced snowpack (**Fig. 3B**). Similarly, drought conditions were observed during the summer of 2017 (**Fig. 3B, 3D**). While seasonal drought indices suggest a generally drier season (**Fig. 3D**), the daily soil moisture index (SMI) values indicate comparatively lower drought intensity in 2017 when juxtaposed with the conditions in 2012 (**Fig. 3C**).

**Figure 3.**
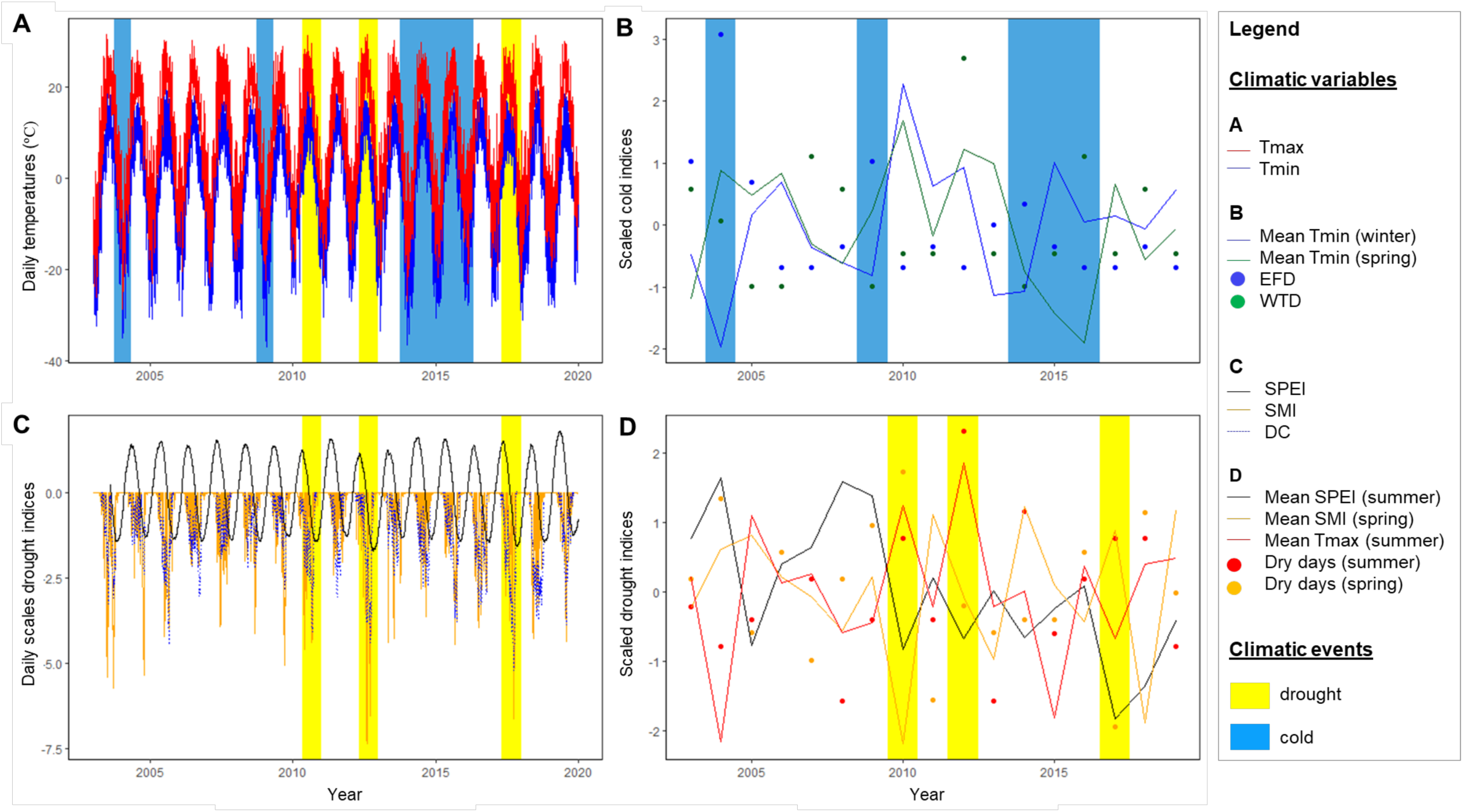
Climatic data for the period 2003-2019 and identification of drought and cold stress periods on the study site. (A) Temporal variation of daily minimum and maximum temperatures. (B) Temporal variation in cold indices, including minimum temperatures (Tmin) in spring (green line) and winter (blue line), the number of frost days (EFD; blue dots), and the number of thaw days (WTD; green dot). (C) Daily temporal variation of the three most relevant drought-indicating climate indices, including soil moisture index (SMI), drought code (DC) and Standardized Precipitation Evapotranspiration Index (SPEI). (D) Annual variation in seasonal drought climate variables, including SMI in spring and summer (orange and red lines, respectively), SPEI (black line), and number of dry days in spring and summer (orange and red dots, respectively). Climatic stress periods identified for the studied timespan are indicated by blue (cold) and yellow (drought) highlighted boxes on the four panels. The complete description of the climatic variables tested in this study can be found in Table 1.

### 3.3. Wood traits connection with climate, and climate-sensitivity traits

The detrended time series of wood parameters displayed significant correlations with both annual and seasonal values of the examined climatic variables (**Fig. 4, Fig. S2**). Earlywood radial growth (ewBAI) exhibited negative associations with previous spring’s late frost risk (LFR) and extreme frost days (EFD) during the preceding winter (**Fig. 4**). Radial growth (BAI and ewBAI) also showed positive correlations with soil moisture during the summer (SMI_Sum) and autumn (SMI_Aut) of the previous year (**Fig. 4A**). Wood density parameters were correlated with the current season’s summer drought (SMI_Sum and SPEI_Sum) and maximum temperatures (TMax_Sum; **Fig. 4B**). Additionally, they were linked to minimal temperatures during the preceding winter (TMin_Win), as well as variables from earlier seasons such as spring soil moisture index (SMI_Spr), maximum summer temperatures (TMax_Sum), and winter thaw days (WTD) from the previous year (**Fig. 4A**). Latewood percentage showed a positive correlation with the maximum temperature of the previous autumn (**Fig. 4A**).

**Figure 4.**
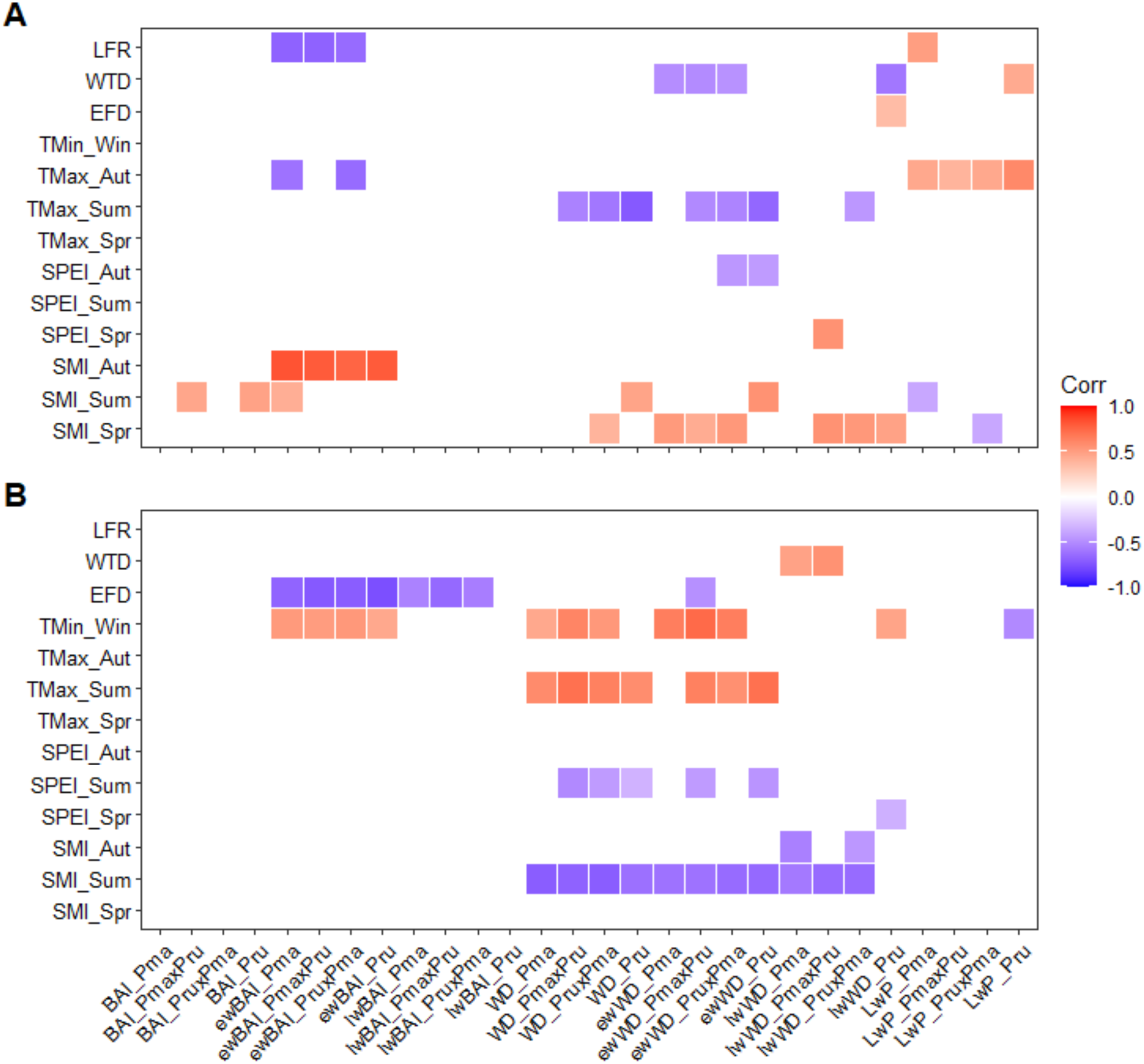
Heatmaps showing the correlations between the wood trait values of each group and the climatic variables tested for the previous year (A) and the current growing year (B). Only significant correlations are shown in color in the heatmaps, positive relationships being indicated in red and negative relationships in blue. Climatic analyses were carried out for radial growth and wood density at whole-ring scale (BAI and WD, respectively), earlywood scale (ewBAI and ewWD, respectively) as well as latewood scale (lwBAI and lwWD, respectively), and also included latewood percentage (LwP) The group names are specified at the end of the wood trait names as follows: *Pma* for *Picea mariana*, *Pru* for *Picea rubens*, *Pma* x *Pru* for the first hybrid type and *Pru* x *Pma* for the second hybrid type. Abbreviations and description of the climatic variables presented here are reported in **Table 1**.

Among the 15 climate-sensitivity traits examined, 10 demonstrated significant group effects, while eight exhibited significant family effects (**Table 3, Table S7**). Six CS traits had consistent genetic effects at the family and group levels (**Table 3**). These CS traits included the sensitivity of earlywood BAI to the preceding year’s autumnal soil moisture, maximum temperature, late frost risk, and minimal winter temperature (ewBAI_SMI_PrevAut, ewBAI_TMax_PrevAut, ewBAI_LFR_Prev, ewBAI_Tmin_Win). They also comprised the CS traits of earlywood WD to the previous summer’s maximum temperature and of latewood WD to soil moisture of the previous spring (ewWD_TMax_PrevSum, lwWD_SMI_PrevSpr).

**Table 3.**
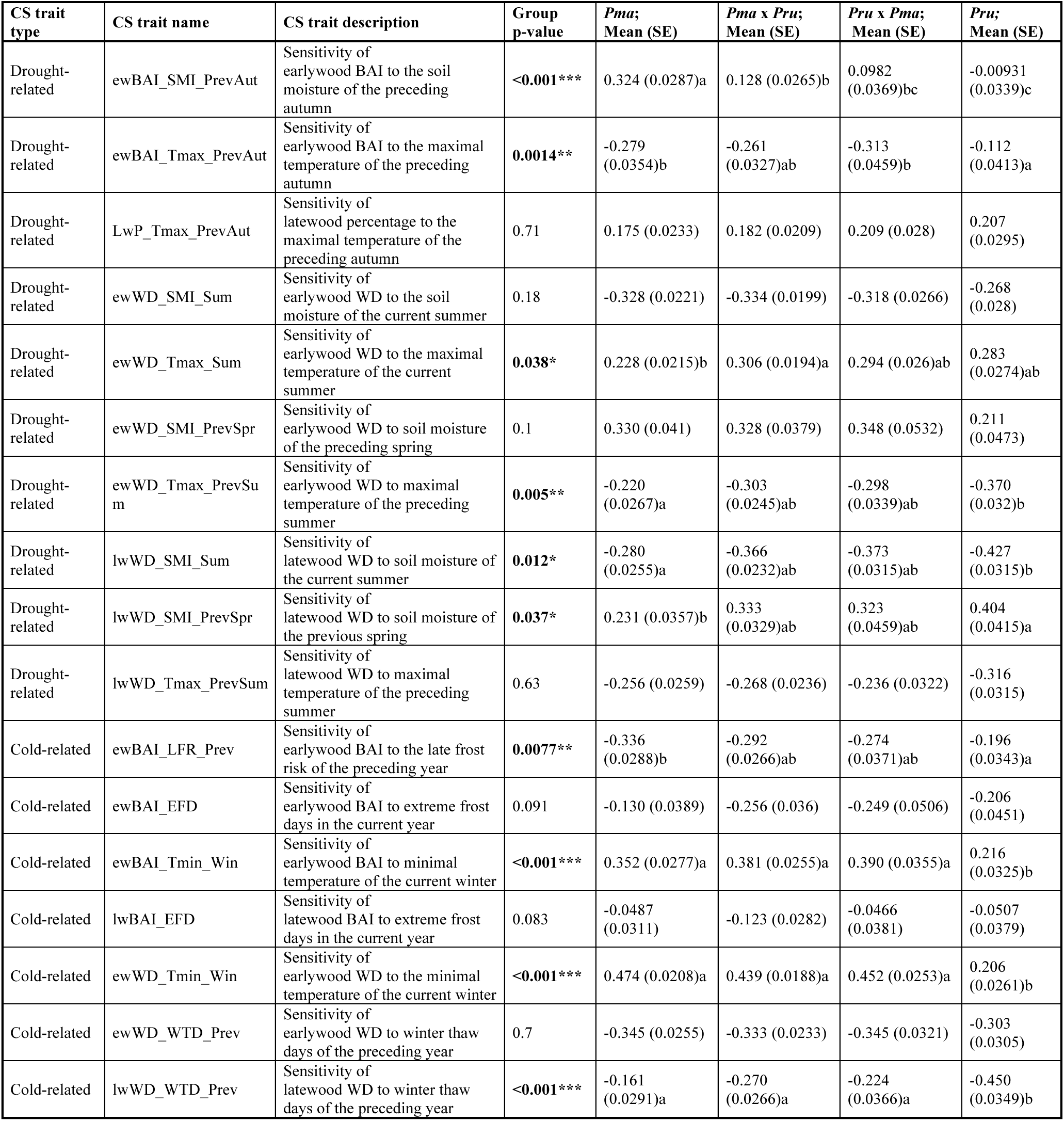
Linear modeling results for the climate-sensitivity traits estimated in this study. Levels of significance for the group effect tested are reported as follows:* *P* < 0.05, ** *P* < 0.01, and *** *P* < 0.001. Whenever the group effect was significant (*P* < 0.05), post-hoc statistical grouping of groups was conducted using the Tukey’s pairwise comparison method with a significance level of 0.05, and significant differences between groups are indicated by different letters. In this table, *Pma x Pru* and *Pru x Pma* represent the two hybrid groups, with *Pma* denoting pure black spruce trees and *Pru* denoting pure red spruce trees.

Significant differences were observed among taxonomic groups in CS traits linked to soil moisture (SMI), maximum temperatures (TMax), extreme winter minimums (EFD, TMin), winter thaws (WTD), and the risk of spring frost (LFR) (**Table 3, Table S7**). Also, no significant differences were observed among groups for the CS traits of latewood percentage to the maximum temperatures of the previous autumn. In comparison to the other three taxonomic groups, red spruce exhibited relatively lower sensitivity to climatic variables in relation with earlywood growth and wood density (ewBAI and ewWD; **Table 3**). The earlywood growth of red spruce was less influenced by maximum temperatures and soil moisture from the preceding autumn (ewBAI_TMax_PrevAut and ewBAI_SMI_PrevAut). Furthermore, red spruce displayed reduced sensitivity of earlywood BAI and wood density to the winter minimum temperatures of the current year (ewBAI_TMin_Win, ewWD_TMin_Win), and to previous year’s late frost risk (ewBAI_LFR_Prev) when compared to black spruce and hybrids. Conversely, regarding earlywood density, red spruce showed higher sensitivity than black spruce to previous summer maximum temperatures (ewWD_TMax_PrevSum), for latewood density in relation to the current summer and the previous spring’s moisture (lwWD_SMI_Sum, lwWD_SMI_PrevSpr), and for winter thaw days of the previous winter (lwWD_WTD_Prev; **Table 3)**.

For two CS traits, the sensitivity of earlywood BAI to the cold and maximum temperatures of the preceding autumn and the sensitivity of latewood density to previous winter thaw days (ewBAI_TMax_PrevAut, lwWD_WTD_Prev) was similar for the two hybrid groups compared to that of black spruce, while climate sensitivity was higher for red spruce. The same trends were observed for the sensitivity of earlywood BAI and earlywood density in response to the minimum temperatures of the current year’s winter (ewBAI_Tmin_Win, ewWD_TMin_Win). For the remaining five CS traits where group differences were found significant (ewBAI_SMI_PrevAut, ewBAI_LFR_Prev, ewWD_TMax_PrevSum, lwWD_SMI_Sum, lwWD_SMI_PrevSpr), hybrids had values intermediate between black spruce and red spruce (**Table 3**).

### 3.4. Response of wood traits to climatic stress periods in red spruce, black spruce, and both hybrid types

Several significant climatic stress periods were observed during the study period (**Fig. 3, Table S3**). Three of these periods met our inclusion criteria (see Material and Methods for details) and were found to be closely linked to changes in five studied wood traits (**Figs. 2 and 3**). In particular, the droughts of 2010 and 2012 (**Table S3**) aligned with increases in wood density (**Fig. 2B**, **2E-F**) and latewood percentage (**Fig. 2G**) in the year of stress occurrence, along with a marked decrease in radial growth in the earlywood and latewood series in the following year or two (**Fig. 2C-D**). During the 2014-2016 period, unusual winter conditions were observed (**Fig. 3B, Table S3**), which were followed by a decline in radial growth in 2016 (**Fig. 2C-D**), an increase in wood density in 2015-2016 (**Fig. 2E-F**), and a peak in latewood percentage in 2016 (**Fig. 2G**).

Significant taxonomic effects were statistically observed across multiple resilience components, covering all assessed wood traits and the three climatic stress periods previously identified using detrended chronologies (**Table 4, Table S8, Table S9)**. Among the tested wood traits, lwBAI displayed the fewest resilience indices significantly associated with a group effect. On the contrary, ewBAI and WD exhibited clear genetic differentiation in their reactions to climatic stress periods, with significant effects from both group and family, being highly significant (*P* < 0.01; **Table 4**).

**Table 4.** Results of linear modeling for two estimated resilience components, Rs and Rc, are provided for the three identified climatic stress periods during the study period. The two resilience components, resistance (Rs) and recovery (Rc), were calculated from detrended data of traits from both earlywood and latewood series of radial growth and wood density. Significance levels for group and family effects are indicated as follows: * *P* < 0.05, ** *P* < 0.01, and *** *P* < 0.001. In cases where the group effect was significant (*P* < 0.05), Tukey’s pairwise comparison method was used for post-hoc statistical grouping of groups with a significance level of 0.05, and significant differences between groups are indicated by different letters. If needed, logarithmic transformations were applied to respect data normality. Trait definitions, ewBAI: earlywood basal area increment; lwBAI: latewood basal area increment; ewWD: earlywood wood density; lwWD: latewood wood density; LwP: latewood percentage.

| Associated climatic stress | Trait | Resilience component | Family effect | Group effect | *Pma*; Mean (SE) | *Pma x Pru*; Mean (SE) | *Pru x Pma*; Mean (SE) | *Pru*; Mean (SE) | Data transformation |
| --- | --- | --- | --- | --- | --- | --- | --- | --- | --- |
| 2010 summer drought | ewBAI | Rs | **0.0057\*\*** | **<0.001\*\*\*** | 0.790 (0.0197)b | 0.820 (0.0181)b | 0.856 (0.0253)ab | 0.923 (0.0225)a | - |
|  |  | Rc | **<0.001\*\*\*** | **0.013\*** | 1.46 (0.0489)a | 1.32 (0.0451)ab | 1.28 (0.0626)ab | 1.25 (0.0547)b | - |
|  | ewWD | Rs | **0.019\*** | **<0.001\*\*\*** | 1.09 (0.00576)a | 1.07 (0.00529)a | 1.08 (0.00758)a | 1.03 (0.00724)b | - |
|  |  | Rc | **<0.001\*\*\*** | **0.59** | 0.947 (0.00656) | 0.950 (0.00605) | 0.945 (0.00872) | 0.958 (0.00782) | - |
|  | lwBAI | Rs | 0.41 | 0.82 | 0.936 (0.047) | 0.953 (0.0436) | 0.977 (0.0619) | 0.908 (0.0544) | log |
|  |  | Rc | 0.5 | **0.0025\*\*** | 1.00 (0.0409)b | 1.10 (0.0402)ab | 1.09 (0.0545)ab | 1.27 (0.061)a | log |
|  | lwWD | Rs | 0.29 | **<0.001\*\*\*** | 0.997 (0.00667)b | 1.02 (0.00607)b | 1.01 (0.00859)b | 1.06 (0.00894)a | - |
|  |  | Rc | 0.5 | 0.26 | 0.986 (0.00558) | 0.974 (0.00506) | 0.974 (0.00711) | 0.969 (0.00773) | - |
|  | LwP | Rs | 0.42 | 0.41 | 1.18 (0.0603) | 1.15 (0.054) | 1.15 (0.0745) | 1.04 (0.0635) | log |
|  |  | Rc | 0.061 | **<0.001\*\*\*** | 0.742 (0.0402)b | 0.861 (0.0426)ab | 0.875 (0.0604)ab | 1.05 (0.0654)a | log |
| 2012 summer drought | ewBAI | Rs | **<0.001\*\*\*** | **0.0019\*\*** | 0.912 (0.0197)b | 0.975 (0.0182)ab | 0.967 (0.0249)ab | 1.02 (0.0222)a | - |
|  |  | Rc | **0.017\*** | **<0.001\*\*\*** | 1.15 (0.0188)a | 1.00 (0.0173)b | 0.986 (0.0236)b | 0.991 (0.0214)b | - |
|  | ewWD | Rs | **<0.001\*\*\*** | **0.032\*** | 1.07 (0.00795)a | 1.05 (0.00732)a | 1.05 (0.0101)a | 1.04 (0.00888)a | - |
|  |  | Rc | 0.11 | **<0.001\*\*\*** | 0.928 (0.00443)b | 0.952 (0.00404)a | 0.954 (0.00553)a | 0.968 (0.00504)a | - |
|  | lwBAI | Rs | 0.5 | **0.005\*\*** | 1.05 (0.0464)a | 0.899 (0.0372)ab | 0.964 (0.0529)ab | 0.834 (0.044)b | log |
|  |  | Rc | 0.094 | 0.21 | 0.979 (0.0553) | 1.05 (0.0551) | 0.884 (0.0629) | 1.03 (0.0674) | log |
|  | lwWD | Rs | 0.056 | 0.24 | 1.05 (0.00771) | 1.06 (0.00705) | 1.07 (0.00966) | 1.07 (0.00875) | - |
|  |  | Rc | 0.26 | **<0.001\*\*\*** | 0.95 (0.0057)a | 0.933 (0.00518)a | 0.93 (0.00704)ab | 0.906 (0.00653)b | - |
|  | LwP | Rs | 0.13 | **<0.001\*\*\*** | 1.27 (0.0705)a | 1.02 (0.0528)ab | 1.08 (0.0755)ab | 0.913 (0.0587)b | log |
|  |  | Rc | 0.5 | **<0.001\*\*\*** | 0.716 (0.0311)c | 0.835 (0.0335)ab | 0.791 (0.042)bc | 0.965 (0.0494)a | log |
| Cold conditions from 2014 to 2016 | ewBAI | Rs | **0.019\*** | 0.097 | 0.881 (0.0202) | 0.943 (0.0188) | 0.887 (0.0253) | 0.889 (0.0228) | - |
|  |  | Rc | **0.0036\*\*** | **<0.001\*\*\*** | 0.963 (0.0268)b | 1.03 (0.0249)b | 1.06 (0.0336)ab | 1.16 (0.03)a | - |
|  | ewWD | Rs | **<0.001\*\*\*** | **<0.001\*\*\*** | 1.06 (0.00616)a | 1.03 (0.00574)b | 1.03 (0.00775)ab | 0.998 (0.00689)c | - |
|  |  | Rc | **0.0062\*\*** | **<0.001\*\*\*** | 0.959 (0.00458)c | 0.977 (0.00426)b | 0.982 (0.00572)b | 1.00 (0.00511)a | - |
|  | lwBAI | Rs | 0.3 | 0.5 | 0.899 (0.0396) | 0.891 (0.0363) | 0.977 (0.0532) | 0.961 (0.0491) | log |
|  |  | Rc | 0.32 | 0.29 | 1.22 (0.0551) | 1.14 (0.0477) | 1.15 (0.0647) | 1.07 (0.0564) | log |
|  | lwWD | Rs | **0.0043\*\*** | **0.0053\*\*** | 1.02 (0.00733)b | 1.04 (0.00681)ab | 1.04 (0.00915)ab | 1.07 (0.00815)a | - |
|  |  | Rc | **0.019\*** | **<0.001\*\*\*** | 1.01 (0.0063)a | 1.00 (0.00584)a | 1.00 (0.00782)ab | 0.974 (0.00702)b | - |
|  | LwP | Rs | 0.092 | **0.026\*** | 1.46 (0.0751)a | 1.25 (0.0604)a | 1.31 (0.084)a | 1.20 (0.07)a | log |
|  |  | Rc | 0.5 | 0.15 | 0.879 (0.0371) | 0.990 (0.039) | 0.990 (0.0507) | 0.941 (0.0459) | log |

The responses of wood density and growth to the 2010 drought period were variable among groups (**Table 4**). Group effects were notably pronounced for the Resistance indices (Rs) of earlywood series for both BAI and WD, as well as for latewood WD. Recovery for latewood BAI and LwP differed among groups, with red spruce showing higher recovery values compared to the other three groups. The 2012 drought also had a differential impact on LwP, earlywood BAI and earlywood WD of the four groups. Group differences were particularly pronounced for the recovery index (*P* < 0.001; **Table 4**). In the latewood series, a significant differentiation among groups was observed for BAI resistance and WD recovery. Those two traits had significantly lower values for red spruce, compared to other groups.

In response to the 2014-2016 winter stress period, groups exhibited different responses in the recovery of earlywood growth (**Table 4**). Highly significant effects were observed at both taxonomic and family levels during the response and recovery phases for wood density during the two growth season phases (0.0035 < *P* < 0.001 for ewWD and lwWD; **Table 4**).

For the majority of resilience traits that differed among groups, the two hybrid groups exhibited intermediate values and were statistically distinguishable from one or both of the pure species groups (**Table 4**), but not from one another.

### 3.5. Phenotypic correlations of growth, wood anatomy and climate-related traits

We observed few significant phenotypic correlations between climate-sensitivity traits and components of resilience, and dendrometric and average wood anatomy traits across groups (**Fig. 5**). The sensitivity of BAI to late frost risk and extreme frost days (ewBAI_Prev_LFR, ewBAI_EFD and lwBAI_EFD) and the sensitivity of earlywood WD to previous summer’s maximum temperature (ewWD_Prev_Tmax_Sum) exhibited negative correlations with cumulative growth variables (H_20, and DBH_20). Conversely, the sensitivity of earlywood WD to minimum temperature, summer’s maximum temperature and previous spring’s soil moisture (ewWD_TNn_Win, ewWD_Tmax_Sum and ewWD_Prev_SMI_Spr) showed positive correlations with H_20 and DBH_20, respectively. When analyzed separately for different groups to identify divergent phenotypic correlation trends, the negative correlation between earlywood BAI sensitivity to previous LFR and growth traits remained significant for black and red spruce, but not for hybrids (**Fig. 5, Fig. S3**). Other associations between CS traits and cumulative growth did not persist at the group level (**Fig. S3**).

**Figure 5.**
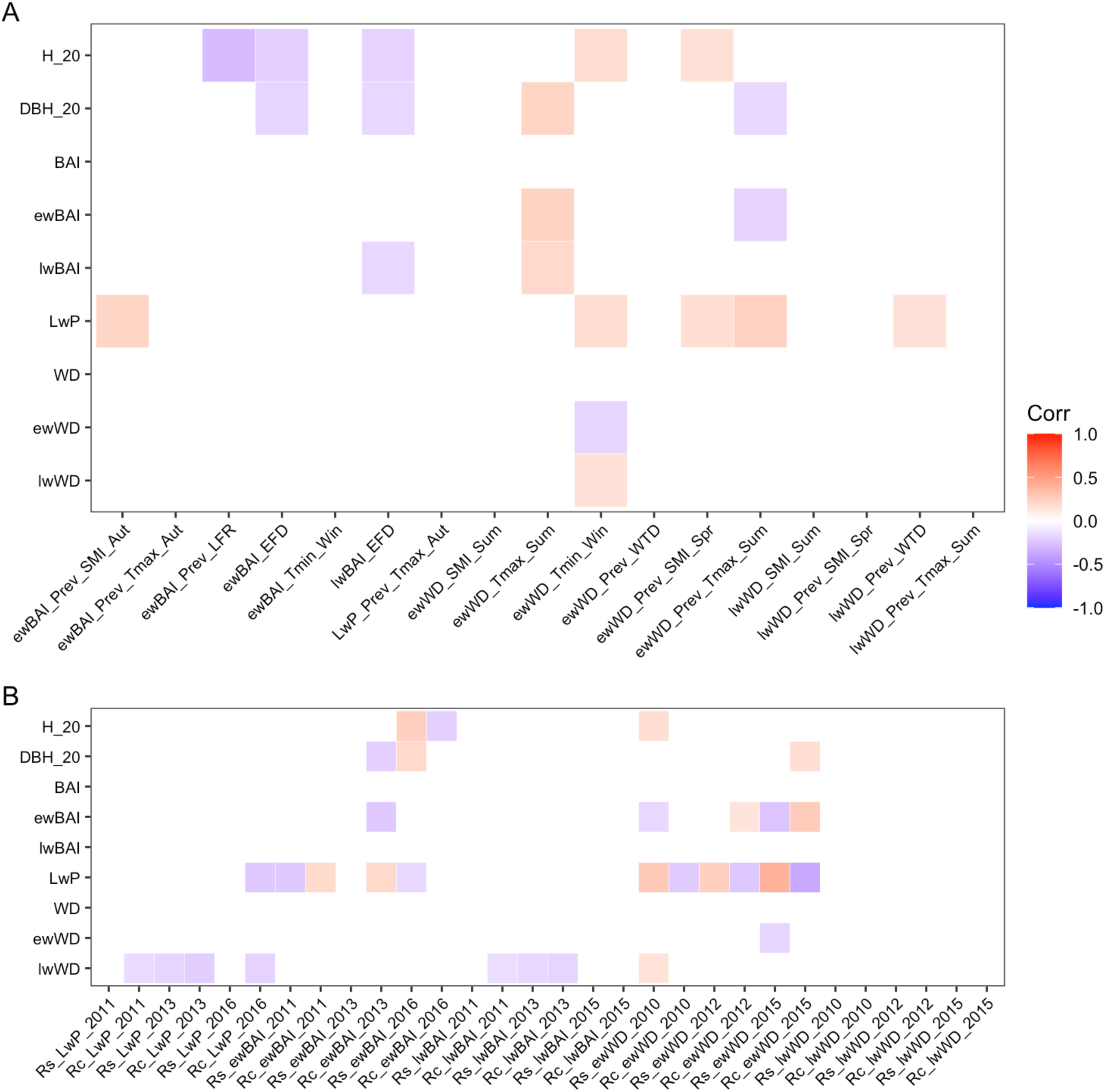
Heatmaps of all-groups phenotypic correlations among the traits analyzed. Pearson correlations were estimated between dendrometric measurements, average annual traits of earlywood, latewood, and total ring series for both wood density and radial growth, and climate-related traits: (**A**) climate-sensitivity traits (see Table 3 for detailed trait descriptions), and (**B**) resilience components (see Table 4 for detailed trait descriptions). Significantly correlated traits after Holm’s adjustment of p-values are highlighted in color, with positive associations shown in red and negative associations in blue.

Across groups, positive correlations were observed between three resilience indices (Rs_ewWD_2010, Rc_ewWD_2015, Rs_ewBAI_2016) and at least one cumulative long-term growth trait (H_20 or DBH_20; **Fig. 5**). When analyzed separately by group to identify divergent phenotypic correlation trends, the resistance of earlywood BAI to cold conditions in 2016 continued to be significantly and positively correlated with growth in all but the *Pru* x *Pma* hybrid group (**Fig. 5**). On the other hand, negative correlations were observed between cumulative growth and the recovery of earlywood BAI in 2013, as well as the recovery of earlywood BAI in 2016 (**Fig. 5**). These negative correlations also did not persist at the group level (**Fig. S3**).

## 4. Discussion

This research work provides valuable and rare insights into the juvenile radial growth patterns and the tree-ring responses of two ecologically differentiated conifer sister species and their F_1_ hybrids to various episodic drought and cold periods under northern mid-latitude climate. The increasing occurrence of climatic stress periods in forests of northern mid-latitudes, driven by climate change and its increasing instability, and the impact of natural hybridization on evolutionary processes represent pivotal factors influencing adaptation traits in many tree species. The specific experimental design established in 1994 allowed us to rigorously compare the two interspecific hybrid types with intraspecific progenies having identical parental origins. In essence, the results of the progeny tests presented here facilitate a formal comparative evaluation of the growth, climate sensitivity, and climate resilience of red spruce, black spruce, and both F_1_ hybrid types.

### 4.1. Differential wood density and growth patterns among groups

In this study, the radial pattern of wood density showed a rapid decrease near the pith followed by a gradual increase towards the bark **(Fig. 2B)**, as previously observed in black spruce (Alteyrac et al., 2005) and white spruce (Soro et al., 2022). F_1_ hybrids displayed intermediate to lower wood density during the second period **(Table 2)**. These wood density rankings, coupled with an intermediate latewood percentage, also placed them within an intermediate to lower range for overall ring wood density. Thus, our results mostly concur with previous findings in *Pinus*, *Eucalyptus* and *Larix* interspecific crosses, where F_1_ hybrids inherited wood density and latewood percentage levels intermediate to those of their parents (Rockwood et al., 1991, Retief & Stanger, 2009, Marchal et al., 2017). This should not be interpreted as a surprising result, given that wood traits in spruces appear to be controlled mostly by additive genetic effects with little dominance effects, when compared to growth traits (Nadeau et al., 2023). In addition, the fact that some wood density values for hybrids were equivalent to or slightly lower than the lowest values recorded for pure species may be due to the typically high radial growth rate of hybrids, as growth and wood density are usually negatively correlated in spruces (*e.g.* Zhang & Morgenstern, 1995).

The radial growth patterns in BAI initially followed an exponential trend but later stabilized **(Fig. 2A)**, in accordance with previous growth patterns observed in spruce species (Koubaa et al., 2005; Lenz et al., 2010). The notable differences in growth patterns noted between black and red spruce, such as black spruce showing higher early growth and red spruce exhibiting increased growth after 13 years **(Fig. 2A**, **Table 2)**, also align well with established silvicultural knowledge about their differing life history strategies. Black spruce is indeed ecologically more competitive at an early age than red spruce, and red spruce is more tolerant of shade, an aptitude possibly favorable in later stages as the tree crowns started closing (Beaulieu et al., 1989; Viereck & Johnston, 1990; Major et al., 2003a; Major et al., 2005; Major et al., 2015). F_1_ hybrids between black and red spruce showed superior cumulative growth trends than the pure species **(Table 2, Table S5)**, likely due to the combination of favorable growth characteristics from both parent species, in a pattern suggesting preponderance of non-additive genetic effects. Effectively, both hybrid types displayed high early growth rate reminiscent of black spruce, and high later growth reminiscent of red spruce, for both earlywood and latewood components, and without reciprocal effects between hybrid crossing directions. Similar dominance effects were noted for morphological traits (Perron & Bousquet 1997). These observations contrast with previous assessments of hybrids of various putative introgressive hybridization degrees at the seedling stage, which indicated slower growth of putative hybrids compared to the pure species (Manley & Ledig, 1979), or at best an intermediate growth profile (Major et al., 2005). Older putative hybrids had also been reported to exhibit growth levels intermediate between black spruce and red spruce (Johnsen et al., 1998; Major et al., 2003b), or at least superior to that of red spruce (Morgenstern et al., 1981).

The lack of concordance between our growth observations and those of previous studies in red and black spruce putative hybrids (Manley & Ledig, 1979; Johnsen et al., 1998; Major et al., 2005) may be attributed to the molecular validation of the parent trees used in our crossbreeding scheme, ascertaining the pure parental or F_1_ hybrid status of the plant material tested. Hybrid vigor has been observed in various traits through many examples of F_1_ interspecific crosses between tree species; however, its expression may be environment-specific or confined to particular crosses owing to the considerable genetic variability usually characterizing largely undomesticated forest tree species (Blada, 2004; Dieters & Brawner, 2007; Volker et al., 2008). Hybrid vigor may diminish in subsequent generations or may not occur if the parents used are already introgressed hybrids at various levels (Rieseberg & Carney, 1998). The unconfirmed parentage of hybrids in prior studies precludes their classification as pure F_1_, given that hybrid indices solely based on morphological traits and traditionally utilized for parental identification were shown to be unreliable in predicting taxonomic identity, especially for introgressed hybrids (Perron & Bousquet, 1997). On a more fundamental level, the superiority of cumulative growth in F_1_ hybrids also tells that fitness related to competitive ability in intermediate environments does not appear to be a barrier to hybridization in the red and black spruce hybrid zone, as opposed to what was reported in some other tree hybridization cases such as *Populus* hybrids in Europe (Christe et al., 2016). It is nonetheless important to mention that our study only examined a single common garden environment located in the zone of contact, cautioning against generalizing the superior growth of F_1_ hybrids to different environments more typical of the allopatric zones of each species. Indeed, previous reports indicated a maximum growth performance of natural hybrids and introgressed trees in environments intermediate to those typical of parental species (De La Torre et al., 2014, Hamilton et al., 2015), such as that tested in our study. Nevertheless, in the context of climate change and environmentally shifting growth conditions, such intermediate environmental conditions may become the norm in new areas where the fitness of hybrids could be superior to that of the parental species, especially if local adaptation is not optimal and thus, parental species suffer locally from maladaptation (Andalo et al., 2005).

### 4.2. Impact of drought and heat on wood traits during tree lifespan

Low soil moisture and high temperatures had immediate, current-year impacts on wood density **(Fig. 4)**, which is largely consistent with previous findings in spruce species (van der Maaten-Theunissen et al., 2013, Soro et al., 2023b). Temperature and water availability have been shown to impact the processes of cell expansion and differentiation, which subsequently determine the final tracheid dimensions and cell wall thickness, ultimately influencing wood density (Balducci et al., 2016; Cuny & Rathgeber, 2016). Under drought conditions, developing xylem cells experience a loss of turgor pressure due to a reduced concentration of osmotically active solutes in the cambial zone, coupled with decreased fixed carbon allocation to xylem cells, leading to a reduction in tracheid diameter and thicker cell walls (Lautner, 2013). Drought-induced changes in cell wall thickness were previously observed in both earlywood and latewood tracheids of Norway spruce (Jyske et al., 2010). Similarly, young white spruce trees under drought treatments exhibited increased wood density and reduced lumen diameter (Soro et al., 2023a, 2023b). The increased wood density that we observed in response to drought (**Fig. 4**) could thus likely be the result of thickened cell walls, whether or not accompanied by a reduction in tracheid diameter.

Radial growth exhibited positive correlations with soil moisture and negative correlations with maximum temperatures during the preceding autumn, implying that the drought of the previous year had a detrimental impact on the current year’s growing season (**Fig. 4**). Accordingly, maximum temperatures of the preceding autumn correlated positively with latewood percentage for three out of four groups **(Fig. 4)**. Prior research has underscored the detrimental impact of drought and heat on wood growth in *P. mariana* and *P. rubens* (e.g. Drobyshev et al., 2013; Yetter et al., 2021). A carry-over or lag effect such as observed herein, where trees exhibited reduced growth in the subsequent year, is typically due to diminished carbohydrate production during drought periods (Bréda et al., 2006). In conifers, the formation of earlywood is heavily dependent on photosynthates from the preceding year, and in cases of secondary growth, there is a potential mobilization of carbon reserves stored over several years, if required (Gessler et al., 2009; Kuptz et al., 2011).

The four groups exhibited variations in climate sensitivity. Compared to red spruce, the earlywood radial growth in black spruce was more responsive to soil moisture availability during the preceding growing season and autumnal maximal temperatures **(Table 3)**. Indeed, this species is recognized for its ability to promptly respond to enhanced water availability through high plasticity in cell production, as observed under severe drought conditions (Balducci et al., 2013). Furthermore, in contrast to red spruce, black spruce exhibited less adverse effects on earlywood density due to the maximum temperature of the preceding summer. Additionally, latewood density showed reduced sensitivity to soil moisture from the previous spring and the current summer (**Table 3**), likely highlighting the ability of black spruce to maintain more stable wood density regardless of moisture conditions.

### 4.3. Impact of cold stress on wood traits: a focus on the pure species, black spruce and red spruce

Negative associations were observed between earlywood radial growth and the LFR variable of the preceding year **(Fig. 4)**. LFR is a variable designed to be indicative of the timing of a late spring frost regarding the accumulation of degree-days, and therefore regarding the state of spring bud phenology or shoot elongation. Late spring frost episodes are well known for causing substantial damage to vulnerable expanding shoots and cambial tissues (Schweingruber, 2007; Benomar et al., 2022), with the most commonly damaging form occurring after bud break, when the well characterized susceptibility of plant tissues is high (Bigras & Hébert, 1996). When a spring frost damages opened buds, they typically result in a marked and immediate reduction in growth for the current year (Dittmar et al., 2006) which did not happen in our study. The likelihood of a bud-damaging late spring frost at our common garden site in southeastern Quebec revealed to be low when considering the climatic data available and the late spring bud phenology of black and red spruce (*e.g.* O’Reilly & Parker, 1982). This leads us to look at another form of spring frost episodes, occurring before bud break in a manner similar to thaw-freeze events, which would be more consistent with the decreases observed in subsequent year’s growth that were significantly associated with this climatic variable, and that came without current-year growth reductions **(Fig. 4)**. Indeed, in conifers, the dehardening process initiates earlier than bud break, resulting in the possibility of freezing injury during strong spring thaw-freeze episodes before bud break, as previously evidenced in black spruce and other conifers (Bigras & Hébert, 1996; Man et al., 2009). In line with this and our observations, Man et al. (2021) reported, from artificial freeze tests simulating spring frosts occurring before bud break, the potential for growth reductions only manifested in the subsequent year. Thus, spring thaw-freeze episodes can lead to growth reductions persisting for several years due to limited resources resulting from the loss of mature needles or buds (Man et al., 2009; Man et al., 2013).

Earlywood and latewood radial growth were significantly correlated with lower winter temperature minima or greater numbers of very cold days throughout the four groups **(Fig. 4)**. This finding aligns well with earlier studies of cold-region continental conifers, where the connection between winter temperatures and ring width is commonly observed (Lindholm, 1996; Cook et al., 1998; Rolland et al., 1999; Wang et al., 2005). In addition to affecting radial growth, colder winter temperatures also resulted in a decrease in wood density values **(Fig. 4)**, consistent with previous findings in black spruce in boreal areas (Xiang et al., 2014; Franceschini et al., 2018). Midwinter minimas are more commonly reported to affect tree rings in environments where trees reach their cold hardiness limits (Suvanto et al., 2017; Hänninen, 2016), although winter temperature was identified as a significant predictor of ring growth also in warmer parts of some species’ range (Pederson et al., 2004). Damage through either freezing injury of needles and buds or winter embolism in stem and roots could lead to a diminished photosynthetic capacity or alterations in resource allocation for damage repair, ultimately drastically reducing growth.

The observed increase in wood density associated with the winter thaw days (**Fig. 4**) may be linked to a reduction in tracheid diameter, a characteristic that could offer protection against thaw-freeze embolism in conifers (Pittermann & Sperry, 2003). But, such a potentially adaptive response, to our knowledge, has not been reported previously in spruce species. The negative correlation of earlywood and latewood density with the number of thaw days of the previous year would be consistent with the documented vulnerability of red spruce to winter thaw-freeze episodes, as evidenced from foliar symptoms and the accompanying radial growth reductions (Wilkinson, 1990; DeHayes et al., 2001). Negative effects from warmer winter temperatures could also result from elevated metabolic activity, increased respiration and energy expenditure (Skre & Nes, 1996).

Out of the seven cold-related climate-sensitivity (CS) traits examined in this study, four exhibited significant differences between black and red spruce (**Table 3**). Black spruce showed more pronounced sensitivities of earlywood BAI to the late frost risk index of the preceding year, and of earlywood density and earlywood BAI in relation to extreme winter minimum temperatures, compared to red spruce (**Table 3**). Conversely, red spruce had a greater sensitivity of latewood density to winter thaws of the previous year (lwWD_WTD_Prev; **Table 3**).

When taken individually, only one of these four CS traits (lwWD_WTD_Prev) is consistent with the predictions where red spruce should have shown more sensitivity, being it is well documented to be more vulnerable to cold and thaw-freeze events than black spruce (*e.g.* Wilkinson, 1990; Strimbeck et al., 1995). The higher sensitivity of earlywood growth and wood density in black spruce indeed contrast with preliminary hypotheses previously reported (DeHayes et al., 2001; Strimbeck et al., 2015). Further, some could argue black spruce’s higher earlywood BAI sensitivity to late frost risk could be due spring dehardening processes could have been slightly earlier than red spruce’s given the geographic origin of the parental trees used, however this interpretation is not quite convincing. Some could say that bud break differences exist between the genetic material from the two species used in our study due to intra- and interspecific latitudinal trends (**Table S1**; e.g. Morgenstern, 1978; Bigras & Hébert, 1996; Morgenstern, 1996; Prakash et al., 2022; Mura et al., 2022), and therefore their spring dehardening should also be affected in the same way. The main counterarguments however are that these interspecific differences are definitely not clear (Major et al., 2005; Major et al., 2015), and if they exist in our material they would be of the scale of up to a few days at most (e.g. Morgenstern, 1978; Major et al., 2015). It would also require that, as was discussed earlier in interpreting the general LFR response, the dehardening process in black spruce would get more quickly to a sensitivity level higher than red spruce’s, which is not comparatively documented, but unlikely given that red spruce has generally lower levels of cold hardiness and much faster dehardening than black spruce during late winter thaw-freeze events (Strimbeck et al., 1995; DeHayes et al., 2001).

The contrast of these results when interpreted as vulnerability and compared to other knowledge suggests that some of the observed CS traits differentiation may be better explained by alternative interpretations such as plastic acclimation or avoidance responses. For instance, the increase in earlywood wood density under winter stress could strengthen cellular structure to enhance long-term frost tolerance, as an adaptive strategy to cope with extremely low winter temperatures. A study on white spruce showed that wood density correlated positively with higher cold hardiness at the population level (Sebastian-Azcona et al., 2018), potentially aligning with this alternative interpretation. The true biological implications and underlying mechanisms of these observations nonetheless remain speculative, highlighting the need for further in-depth investigations, such as testing the material on various sites with contrasting degree-day accumulation and winter temperatures, so to better understand the mechanisms of cold acclimatization in both species.

### 4.4. Differential responses among groups to climatic stress periods: exploring tree ring anatomy responses to cold and drought

In our study, both drought periods detected in 2010 and 2012 led to an increase in wood density during the drought year and a decrease in radial growth in the following year **(Fig. 2**, **Table 4)**. Differences in the degree of trait response were observed, with the 2012 drought being more severe and eliciting more marked reactions in latewood density and radial growth compared to the 2010 drought (**Fig. 3C-D**, **Table 4, Table S3**). This disparity may arise from differences in the intensity and timing of drought, as previously shown in tree species (Forner et al., 2018; Gao et al., 2018; Laverdière et al., 2022). Cumulative effects and response trajectories may also have played a role in modulating wood reactions during the following drought episode, as documented in conifers (Anderegg et al., 2020; Gessler et al., 2020). The pronounced differentiation between groups in 2010 for ewBAI and ewWD components (**Table 4**) suggests varied response to drought among taxonomic groups during the stress resistance phase, in contrast with the recovery phase. However, the response of LwP to climatic stress periods was relatively consistent across taxonomic groups (**Table 4**), indicating that this trait does not effectively discriminate among groups in this context. Interestingly, the average LwP positively correlated with drought response traits for ewBAI and ewWD (Fig. 4B), suggesting that a higher latewood proportion may enhance resistance to drought. This interpretation aligns with the concept that the latewood-to-earlywood ratio influences trees’ drought resistance by modifying hydraulic conductivity and water transport security (Rathgeber, 2017; Fan et al., 2018). The strategy of reducing radial growth during dry years while increasing the latewood proportion has been recently proposed as an adaptive acclimatization strategy response to cope with subsequent episodic droughts in larch trees (Zhang et al., 2024). In our study, red spruce exhibited weaker earlywood responses to drought, while black spruce showed weaker latewood responses, highlighting a contrasting acclimative strategy between the two species.

Following cold conditions, distinct differentiation among groups was observed in terms of wood density and earlywood BAI reactivity (**Table 4**), including group-specific physiological variation during the recovery process after stress. These differences could reflect different physiological strategies between the studied groups, which could eventually result in different resilience levels when facing cold stresses. To date, there has been a scarcity of studies that investigated thoroughly the mechanisms responsible for the recovery of growth and physiological functions in evergreen trees, notably addressing cold-induced embolism after stressful cold episodes.

### 4.5. Wood response patterns in black and red spruce: implications for wood phenology

Across all climate-related tree-ring traits (except latewood percentage) and related to both drought and cold responses, we detected a pattern of significant differences in ring zones response between the two pure species that could be explained by diverging cambial phenology. For both radial growth and wood density traits, black spruce showed a general pattern of higher earlywood response, while red spruce showed higher latewood response (**Table 3**, **Table 4**). This observed interspecific differentiation is likely the result of differing wood phenology. It is well known that intraspecific variation in phenology can be linked to adaptations to differing lengths of growing season in black spruce (e.g. Morgenstern, 1978) and red spruce (e.g. Butnor et al., 2019; Prakash et al., 2022), and in many wide-ranging forest tree species including other northern mid-latitude spruces (Morgenstern, 1996; Li et al., 1997), with provenances originating from warmer climates exhibiting later growth cessation. Common garden studies in a few species indicate that such phenological differences also exist in cambial growth in relation to latitude of origin (Vargas-Hernandez et al., 1994; Jayawickrama et al., 1997). Previous Scots pine provenance studies suggested that differences in cambial phenology between southern and northern seed sources can also explain differential latewood sensitivity to climatic variation, as a result of the offset between the climate they’re adapted to and the test site’s climate (Savva et al., 2003; Savva & Vaganov, 2006). This interpretation of the differential reactivity between the two species being due to different cambial phenologies, is further supported by the greater proportion of latewood in black spruce, our more northern group, a result analogous to the studies cited above (Vargas-Hernandez et al., 1994; Jayawickrama et al., 1997; Savva et al., 2003; Savva & Vaganov, 2006).

Both environmental factors and genetic features interact to determine the phenology of wood formation (Zobel and Jett, 1995; Jayawickrama et al., 1997). In this context, it is also likely that differences in wood phenology between black spruce and red spruce would arise from both genetically-determined developmental as well as environmental factors, leading to distinct wood anatomy responses in each species. In conifers, xylogenesis is largely regulated by photoperiod (Mu et al., 2023; Carteni, 2018) but is also influenced by various internal factors. These factors, interacting with photoperiod, can cause temporal delays in the transition between phases of xylogenesis, such as from earlywood to latewood, without necessarily postponing the cessation of latewood formation at the end of the season (Moser et al., 2010).The resulting potential phenological differences in cambial growth during a given growing season could explain the fact that the two species express similar climatic signals, but in different tree-ring zones. One should note that the exact environmental cues, signaling mechanisms regulating wood formation and their interactions with genetic factors are still poorly understood (Deslauriers et al., 2007; Eckes-Shephard et al., 2022), and more in-depth studies will be required to fill this knowledge gap.

Our study highlights the genetic similarity between black spruce and red spruce, drawing parallels to intraspecific differentiation patterns. In our study, the progeny test site (St-Luc, **Fig. 1**) was situated farther north than the average geographic origin of red spruce parents used for intra- and interspecific crosses. This location, characterized by a colder local climate, led to a shorter growing season compared to southern sites that better represent the local adaptation of the red spruce material examined in this study. Previous studies on intraspecific provenance suggest that genetic material exposed to warmer climates may experience increased stress due to maladaptation, often reflected in reduced growth potential or higher vulnerability to drought (e.g., Carter, 1996; Rehfeldt et al., 1999; Isaac-Renton et al., 2018; Andalo et al., 2005). This "climate warming" effect might have contributed to the greater earlywood reactivity observed in black spruce. This aligns only with the hypothesis that a stronger response indicates higher vulnerability, which still requires confirmation for each specific sensitivity or resilience trait.

### 4.6. Resilience and sensitivity in F_1_ hybrids: similarities to black spruce amidst overall intermediate climate traits

The values of climate-response traits of hybrids were in most cases approximately intermediate between those of pure species., along with the absence of reciprocal or maternal effects between the two directions of hybrid crosses (*Pma* x *Pru* versus *Pru* x *Pma*), both expressing similar phenotypic values. This concurs on a general level with the previous observations of Wilkinson (1990), who suggested that putatively introgressed red spruce populations partly inherited black spruce’s resistance to thaw-freeze stress. It is also analogous to reported results from other spruce hybrids, for instance F_1_ obtained from interspecific crosses between interior and Sitka spruce, where cold hardiness was inherited at an intermediate level between parental species (Kolotelo, 1991), thus suggesting an inheritance pattern mostly dependent on additive genetic effects for these climate-response traits. Interspecific hybrids of other tree species (Duncan et al., 1996; Brennan et al., 2021) have also shown similar inheritance patterns of traits related to climate adaptation.

In some instances, hybrids also exhibited trait values that were more closely aligned with those of black spruce, rather than red spruce (four CS traits; **Table 3**). We hypothesize that this significant inclination of certain hybrid traits towards black spruce, the progenitor species of the pair, may result from the prevalence of dominance at certain underlying loci where red spruce carries recessive alleles. This could first be possible because of differential adaptation in the two species. It is indeed likely that, due to red spruce’s isolation from black spruce under milder climate conditions on the eastern coast of U.S.A. at species inception during one of the Pleistocene glaciations (Perron et al., 2000), local adaptation to such mild conditions would have prevailed during many generations, hence contributing to the rapid fixation of favorable alleles in the founding red spruce population. But the prevalence of dominance toward black spruce could also be the result of likely stronger genetic drift effects during the history of red spruce since species inception. Indeed, when the founding red spruce ancestral population derived from black spruce during the Pleistocene era, it likely suffered from multiple severe population bottlenecks compared to black spruce (Perron et al., 2000; Capblancq et al., 2020). It may also have endured more recent bottleneck effects related to reductions in population size due to Holocene warming and altitudinal migration of its populations throughout the Appalachian Mountain range (Jaramillo-Correa et al., 2015; Capblancq et al., 2020). Consequently, it has endured important reductions in effective population sizes, increasing the strength of genetic drift effects and the likelihood of fixation of alleles, including recessive ones that could bear adaptive consequences (Perron et al., 2000; Jaramillo-Correa et al., 2015; Capblancq et al., 2020).

### 4.7. Tree-ring derived traits in relation to the more traditional cumulative growth

Dendroecological analysis applied to genetic tests having gained in popularity recently, there are still too few studies regarding the value of tree-ring traits for assessing fine-scale genetic aspects of sensitivity and adaptive capacity of trees to climate change (Aubin et al., 2016; Moran et al., 2017; Depardieu et al. 2020). Understanding how tree-ring derived traits correlate with the more traditional cumulative growth traits used as proxies for adaptation, or even wood anatomy traits, can help decipher their informativity and the underlying genetic architecture between the traits, ultimately guiding their use in tree improvement research (Laverdière et al., 2022). In our study, only few CS traits and drought resilience components were significantly correlated with the cumulative growth traits depicted by tree height and DBH (**Fig. 5**). This contrasts with previous studies, notably on white spruce, where a positive correlation between resilience or recovery from drought and cumulative tree growth was shown (Depardieu et al., 2020; Laverdière et al., 2022). Furthermore, several wood functional traits, such as wood density, latewood percentage, or tracheid length, were independently shown to correlate with species’ growth responses to drought at both short- and long-term scales (Serra-Maluquer et al., 2023; Soro et al., 2023a, 2023b), and being effective proxies for adaptive traits associated with resilience to drought (Domec & Gartner, 2002; Ruiz Diaz et al., 2014). The divergence of our results from these previous studies highlights that the interrelations between the different types of traits are not universal and may be dependent on context, notably that of the climatic stresses encountered, and the limits imposed by the genetic material in test. In this context, more dendroecological studies will be required to assess potentially adaptive wood anatomical traits related to responses to climatic stress.

## 5. Conclusions

We undertook an original long-term assessment under controlled conditions and presented a thorough comparison of annual growth rate and response to drought and cold conditions among the ecologically-contrasted progenitor-derivative *Picea mariana* and *Picea rubens,* and their F_1_ hybrid progeny obtained after *a priori* validation of parental types using molecular fingerprinting. Xylem anatomical traits revealed similar climate limitations, yet showcasing what appears to be phenologically different strategies and short-term responses in black spruce, red spruce, and their two reciprocal hybrid groups. Our findings underscore the imperative to intensify research on wood formation and the mechanisms regulating the phenological shift of cambium physiology, along with its impact on the earlywood-latewood proportion, to predict forest productivity of naturally introgressing species in the context of global warming and more unstable climate. Future multi-site studies with robust experimental designs for high-quality genetic assessments of functional variability (Perron et al., 2013) should prioritize this hybrid complex by including the site studied here, along with a colder site and a southern site representative of a more meridional climate.Such designs would allow to strengthen our understanding of these species’ physiological responses to climate stresses and to properly evaluate the impacts of natural introgressive hybridization on the adaptive potential of these two species in the context of climate change. Forthcoming research should also prioritize the analysis of quantitative genetic variation in naturally introgressed or complex hybrids and assess the sensitivity, acclimation, and adaptation responses of both natural and pedigree populations to the impacts of climate change. Our results provide a foundation for future common-garden studies and genomic research to better understand the genetic basis of climate adaptation in red and black spruce, their hybrids, and the effects of natural hybridization in the context of climate change.

## Declarations

### Availability of data and materials

The datasets generated and analyzed during the current study are available in the *Hybrids red spruce black spruce climate sensitivity resilience* repository, and can be found here: https://github.com/ereedmetayer/PiceaRubensMarianaF1Hybridization. Main R scripts have been deposited in the github repository, and other R scripts will be available from the corresponding author on reasonable request.

### Author’s contributions

MP, JB, PL, and ERM designed the study. ERM, JB, MP, and CD designed methods. ERM and MP carried out the experiments. ERM, CD, and MP performed the analyses and discussed the results. ERM, CD, and MP wrote the manuscript with contributions and feedback from JB and PL. All authors contributed to the article and approved the submitted version.

## Acknowledgements

We thank the Direction de la recherche forestière of the ministère des Ressources naturelles et des Forêts of Quebec (MRNF) for access to the study site (SLC30498) and their staff for sampling trees used in this study. We especially thank Simon Nadeau for parentage verification and Pier-Luc Faucher (MRNF) who led recent technical aspect for this trial. The authors are also grateful to Philippe Labrie, Eric Dussault, Jean-François Légaré, Ioan Nicolae, Samuel Lapierre, France Gagnon, Jean-Philippe Laverdière, Françoise Pelletier, Rosalie Daigle, Esther Pouliot, and Vincent Seigner (Natural Resources Canada) for their assistance in the laboratory and field. The authors express their gratitude to Rémi Saint-Amand (Natural Resources Canada) for developing new climatic models for the BioSIM version 11 software.

