## Supplementary Materials for "Spruce hybrids show superior cumulative growth and intermediate response to climate anomalies, as compared to their ecologically divergent parental species"

### Supplementary material

### Supplementary figures

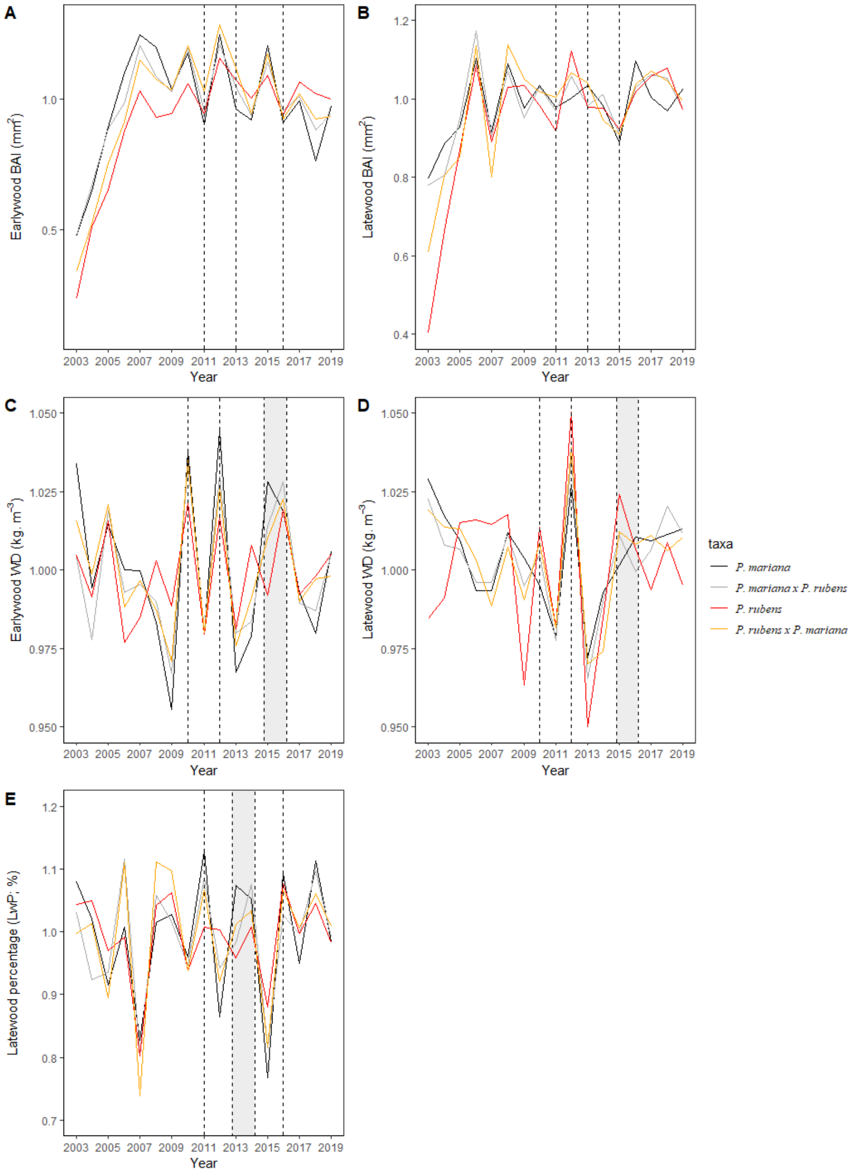

**Figure S1. Inter-annual variations in detrended chronologies for radial growth, wood density and latewood percentage for black spruce, red spruce, and their hybrid groups.**

Average time series by group are presented for earlywood and latewood detrended chronologies of basal area increment (ewBAI and lwBAI, respectively; **A, B**) and wood density (ewWD and lwWD, respectively; **C, D**), as well as for latewood percentage (LwP; **E**). The dotted lines indicate specific years in which substantial variations in earlywood and latewood series were observed following drought and/or cold stress periods experienced by the trees, for the period 2003-2019.

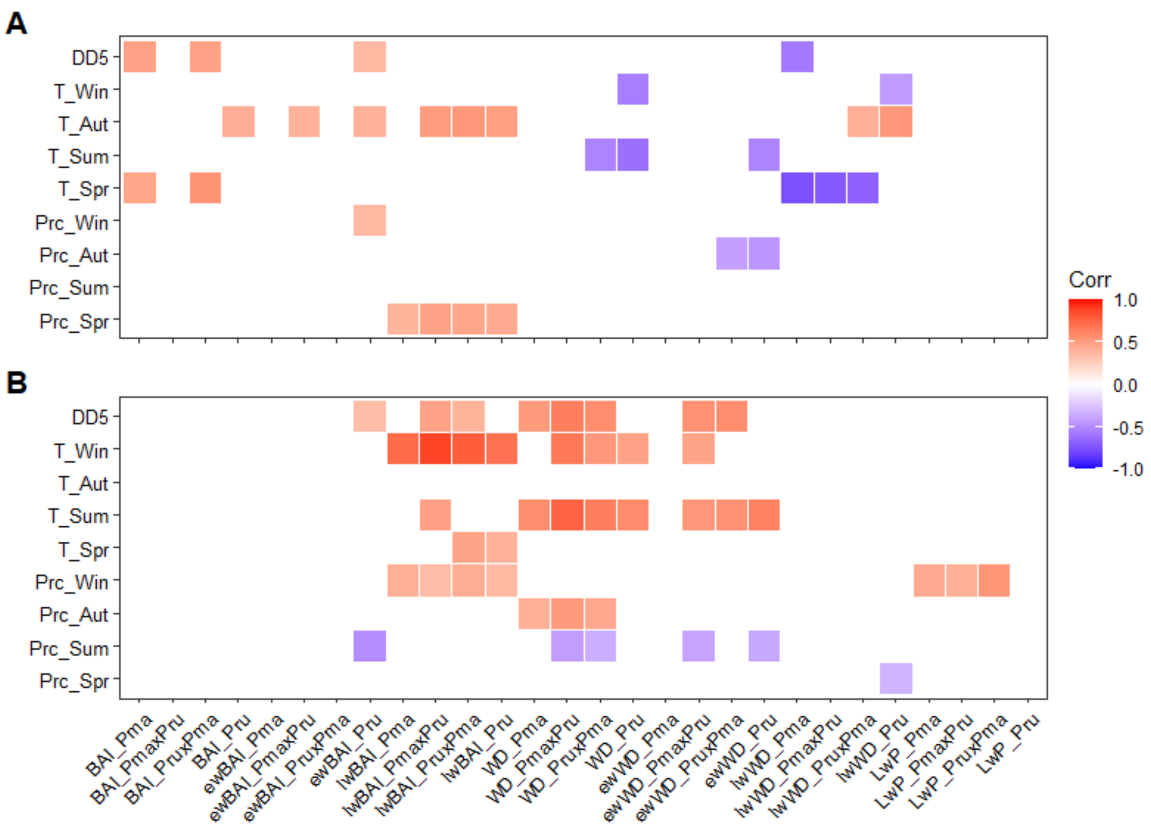

**Figure S2 Heatmaps showing correlations between the wood traits of each group and the climatic variables tested (degree-days, mean temperature and total precipitation) for the previous year (A) and the current growing year (B).**

Only significant correlations are shown in color in the heatmap, positive relationships being indicated in red and negative relationships in blue. Climate correlation analyses were carried out for radial growth and wood density at whole-ring (BAI and WD, respectively), earlywood (ewBAI and ewWD, respectively), as well as latewood scales (lwBAI and lwWD, respectively). The group names are specified at the end of the wood trait names as follows: *Pma* for *Picea mariana*, *Pru* for *Picea rubens*, *Pma* x *Pru* for the first hybrid group and *Pru* x *Pma* for the second hybrid group. Abbreviations and description of the climatic variables are as follows: DD5, growing degree-days (base 5°C); T_season and Prcp_season are mean temperature and total precipitation for a given season, respectively (Win: winter; Aut: autumn; Sum: summer; Spr: spring).

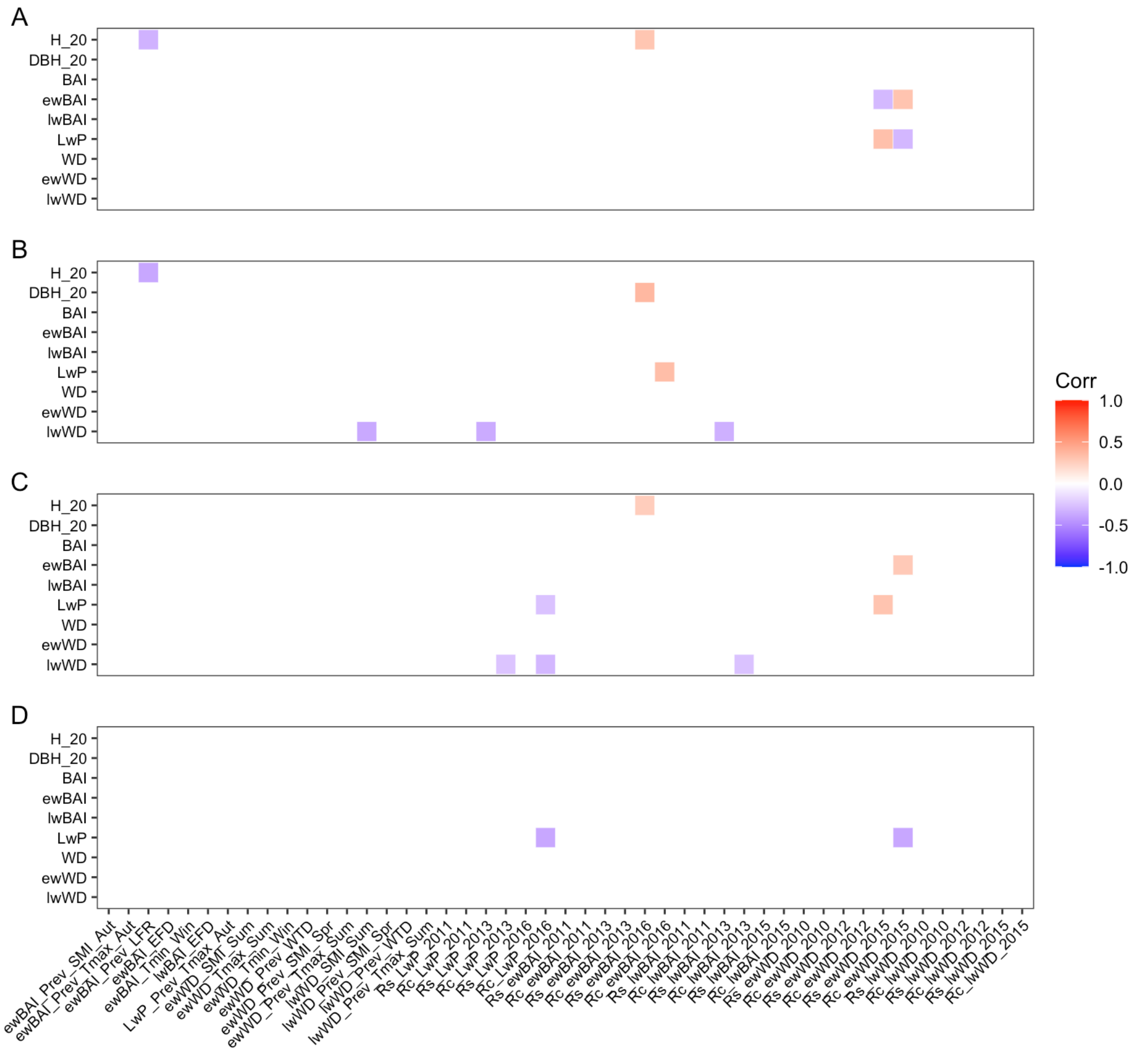

**Figure S3. Group-level phenotypic correlations among the studied traits.**

Pearson correlations were estimated to explore the relationships between dendrometric measurements, average annual traits of earlywood, latewood, and total wood series, both in terms of wood density and radial growth with climate-related traits, including climate sensitivity and resilience components, for the four groups: A) *Picea mariana*, B) *Picea rubens*, C) *P. mariana* x *P. rubens*, and D) *P. rubens* x *P. mariana* hybrids. Traits that exhibited significant correlations, following Holm's adjustment of p-values, are highlighted in color. Positive associations are represented in red, while negative associations are depicted in blue.

### Supplementary tables

**Table S1. Detailed information on the species and geographic origins of the parent trees of the studied families obtained after crosses.**

The maternal geographical origins are shown through the inclusion of provenance names and corresponding geographic coordinates. The pollen-donor species and the recipient mother species for each examined family are also reported.

| Family name | Mother species | Provenance name | Latitude (°) | Longitude (°) | Elevation (m) | Pollen-donor species | Group |
|---|---|---|---|---|---|---|---|
| 20901 | *P. mariana* | Parc Mistassini, Abitibi-Est | 50.45 | -73.63 | 365 | *P. rubens* | *Pma x Pru* |
| 20902 | *P. mariana* | Chûte-aux-galets, Chicoutimi | 48.62 | -71.13 | 185 | *P. rubens* | *Pma x Pru* |
| 20903 | *P. mariana* | Manouane, Laviolette | 47.80 | -74.13 | 460 | *P. rubens* | *Pma x Pru* |
| 20904 | *P. mariana* | Manouane, Laviolette | 47.80 | -74.13 | 460 | *P. rubens* | *Pma x Pru* |
| 20905 | *P. mariana* | R. Tweedie, Northumberland, N.-B. | 46.82 | -65.15 | 60 | *P. rubens* | *Pma x Pru* |
| 20906 | *P. rubens* | October Mts., Mass. | 42.37 | -73.25 | 549 | *P. mariana* | *Pru x Pma* |
| 20907 | *P. rubens* | Essex County, N.Y. | 44.42 | -73.67 | 610 | *P. mariana* | *Pru x Pma* |
| 20908 | *P. rubens* | Essex County, N.Y. | 44.42 | -73.67 | 610 | *P. mariana* | *Pru x Pma* |
| 20909 | *P. rubens* | Saint Charles de Mandeville, P.Q. | 46.50 | -75.33 | 210 | *P. mariana* | *Pru x Pma* |
| 20910 | *P. mariana* | Parc Mistassini, Abitibi-Est | 50.45 | -73.63 | 365 | *P. mariana* | *Pma* |
| 20911 | *P. mariana* | Manicouagan 5, Saguenay | 50.67 | -68.78 | 430 | *P. mariana* | *Pma* |
| 20912 | *P. mariana* | Chûte-aux-galets, Chicoutimi | 48.62 | -71.13 | 185 | *P. mariana* | *Pma* |
| 20913 | *P. mariana* | R. Tweedie, Northumberland, N.-B. | 46.82 | -65.15 | 60 | *P. mariana* | *Pma* |
| 20914 | *P. mariana* | Timmins, Ogden, Ontario | 48.53 | -81.42 | 305 | *P. mariana* | *Pma* |
| 20915 | *P. mariana* | Ipsala, Ontario | 49.00 | -90.45 | 475 | *P. mariana* | *Pma* |
| 20916 | *P. rubens* | Essex County, N.Y. | 44.42 | -73.67 | 610 | *P. rubens* | *Pru* |
| 20917 | *P. rubens* | Amherst, Maine. | 44.90 | -68.38 | 140 | *P. rubens* | *Pru* |
| 20918 | *P. rubens* | Andorra Forest, N.H. | 43.08 | -72.12 | 518 | *P. rubens* | *Pru* |
| 20919 | *P. rubens* | Valcartier Forest Exp. Sta., P.Q. | 46.92 | -71.55 | 274 | *P. rubens* | *Pru* |
| 20920 | *P. rubens* | Saint Charles de Mandeville, P.Q. | 46.50 | -75.33 | 210 | *P. rubens* | *Pru* |
| 20921 | *P. mariana* | Manicouagan 5, Saguenay | 50.67 | -68.78 | 430 | *P. rubens* | *Pma x Pru* |
| 20922 | *P. mariana* | Chûte-aux-galets, Chicoutimi | 48.62 | -71.13 | 185 | *P. rubens* | *Pma x Pru* |

**Table S2. Description and abbreviations of the climatic variables tested in this study.**

| **Variable** | **Definition** | **Comments** |
|---|---|---|
| Variables that were included in the tree-ring stress analysis. | | |
| LFR | Late frost risk; Degree days above 5°C cumulated at time of last spring frost below 0°C. | Spring phenology being strongly associated with cumulative degree-days, more degree-days accumulated at the moment of the last spring frost informs on the risk of frost damage. |
| WTD | Number of thaw days; Number of days where minimum temperature is above 0°C, between December 15$^{th}$ and March 15$^{th}$ of the previous winter. | Winter thaw events with subsequent freezing temperatures have been documented to induce frost damage in red spruce. |
| EFD | Extreme frost days; Number of days with minimum temperature below -30°C, from November to march of the previous winter. | Extreme cold days can cause frost damage to insufficiently cold hardy trees. |
| Tmin | Absolute minimal monthly or seasonal temperature. | Extreme cold days can cause frost damage to insufficiently cold hardy trees |
| Tmax | Absolute maximal monthly or seasonal temperature. | Extremely warm days can cause heat stress. |
| SPEI | Standardized Precipitation Evapotranspiration Index; Mean monthly or seasonal standardized precipitation evapotranspiration index using the Hargreaves method. | SPEI tells about drought severity, informing on drought stress (higher SPEI = more severe drought). |
| SMI | Soil moisture index; Mean monthly or seasonal soil moisture index. | SMI tells about drought severity, informing on drought stress (lower SMI = more severe drought). |
| The following variables were included as primary variables that could underly more stress relevant metrics, or predispose to other stresses, and as complementary information. | | |
| A treeclim correlation analysis of these variables is included as Figure S2. | | |
| DD5 | Summer cumulative degree-days above 5°C. | Summer degree-days are an important indicator of time and heat available to complete the annual growth cycle, thereby influencing time to harden off and to build up energy reserves. |
| T | Mean monthly or seasonal temperature. | Mean temperature is a primary climate variable, with an important impact on cold, heat and drought stress. |
| Prc | Total monthly or seasonal precipitation. | Total precipitation is a primary climate variable, with an important impact on drought stress. |
| The following variables were discarded from further analysis after either not appearing related to chronologies, being strongly collinear with other relevant variables, or being the less relevant variable of a group reflecting the same type of stress (relevance determined from a combination of theoretical basis, and a preliminary PCA analysis). | | |

| | | |
|---|---|---|
| Consecutivedayswithoutf rost | Number of consecutive days without frost from spring to autumn (i.e. the length of the frost free period). | Number of days without frost during the summer is an indicator of time available to complete the annual growth cycle, thereby influencing time to harden off and to build energy reserves. |
| CSDI | Cold spell duration index; Monthly or seasonal index, calculated from the count of days where the minimum temperature is below the 10th percentile for 6 consecutive days or more, from Climdex. | Extreme cold days can cause frost damage to insufficiently cold hardy trees. |
| SDII | Simple precipitation intensity index; Monthly or seasonal index, calculated form the precipitation amounts and the count of wet days, from Climdex. | Indicative of precipitation distribution, could have been relevant for drought stress. |
| Seasonal_Minus20 | Number of days with minimum temperature below -20°C, from November to March of the previous winter. | Extreme cold days can cause frost damage to insufficiently cold hardy trees. |
| FWI_DC | Drought code; Mean monthly or seasonal drought code, from the Canadian Forest Fire Weather Index System (FWI). | DC tells about drought severity, informing on drought stress (higher DC = more severe drought). |
| NbThaw | Number of thaw events; Number of events (consecutive days) where minimum temperature is above 0°C, between December 15th and March 15th of the previous winter. | Number of events. Winter thaw events with subsequent freezing temperatures have been documented to induce frost damage in red spruce. Similar to WTD, but on a number of events basis. |
| LastSub0FrostJulDay | Julian date of the last frost below 0°C. | Indicative of severe late spring frosts, with regard only to julian date. |
| FD | Frost days; Monthly or seasonal count of days where temperature gets below 0°C, from Climdex. | Number of frost days per month could have indicated early winters or late winters, associated with a reduction in growing season time available. |

**Table S3. Characteristics of the climatic stress periods detected.**

The attributes of the three identified climatic stress periods are detailed in terms of their timing, duration, and stress intensity. The duration of the drought episodes was determined based on SMI values (SMI values less than 95 per cent). The corresponding drought code (DC), minimal or maximal temperatures (Tmin and Tmax, respectively) values for these periods are reported.

| **Type** | **Year** | **Season** | **Time span** | **Number of days (consecutive or punctual)** | **Drought intensity (SMI values)** | **Drought intensity (DC values)** | **T (Tmax and Tmin) values** |
|---|---|---|---|---|---|---|---|
| Drought | 2010 | Spring | May 21st – May 31st | 10 consecutive | 90.2 - 94.9 | 72.1 – 118.6 | Tmax : 18.2 - 28.3 |
| Drought | 2010 | Summer | June 18st - June 22st | 4 consecutive | 93.4 - 94.4 | 83.0 – 127.0 | Tmax : 25.4 - 27.7 |
| Drought | 2010 | Summer | August 28st - September 6st | 9 consecutive | 90.6 - 94.6 | 173.5 - 217.0 | Tmax : 13.6 - 29.6 |
| Drought | 2012 | Summer | June 20st - June 24st | 4 consecutive | 93.5 - 94.3 | 57.1 - 84.9 | Tmax : 20.5 - 27.2 |
| Drought | 2012 | Summer | July 12st - July 16st | 4 consecutive | 92.6 - 94.6 | 76.8 – 100.5 | Tmax : 24.5 - 30.8 |
| Drought | 2012 | Summer | July 21st - August 9st | 19 consecutive | 84.5 - 94.1 | 103.4 – 203.1 | Tmax : 18.2 - 28.3 |
| Cold (Tmin < -20) | 2013-14 | Winter | November 30th- March 26th | 49 punctual | na | Na | Tmin: -37.1 |
| Late cold | 2014 | Spring | March 23rd-26th, April 16th-18th | 7 punctual | na | Na | Tmin: -25.8, -13.2 |
| Thaw-freeze | 2014-15 | Winter | December 24th-January 10th | 18 days duration (6 thaw days) | na | Na | Tmax: 1.7 - 4.8; Tmin: -12.5 - -31.1 |
| Cold (Tmin < -20) | 2014-15 | Winter | December 30th- April 6th | 46 punctual | na | Na | Tmin: -31.1 |
| Late cold | 2015 | Spring | March 22nd-24th, April 5th-6th | 2 punctual | na | Na | Tmin: -19.2, -21.4 |
| Thaw-freeze | 2015-16 | Winter | December 23rd- 30th | 8 days duration (4 thaw days) | na | Na | Tmax: 0.8 - 10.5; Tmin: -13.9 - -18.1 |
| Thaw-freeze | 2016 | Winter | January 31st- February 15th | 16 days duration (4 thaw days) | na | Na | Tmax: 3.6 - 5.9; Tmin: -9.2 - - 28.2 |
| Late cold | 2016 | Spring | April 4th - 6th, 10th-11th, 24th-May 1st | 13 punctual | na | Na | Tmin: -19.0, -13.3, -10.7 |

**Table S4. Characteristics of the resilience component indices.**

The year of the wood trait's peak value, along with the duration of both pre-stress and post-stress periods are reported. For each trait analyzed, the resulting names of the three resilience components (Rs: resistance; Rc: recovery; Rl: resilience) are reported.

| Trait | Year pic | Pre-stress period | Post-stress period | Name of the resilience components (Rs, resistance; Rc, recovery; Rl, resilience) |
|---|---|---|---|---|
| ewBAI | 2011 | 2010 (one year) | 2012 (one year) | Rs_ewBAI_2010; Rc_ewBAI_2010; Rl_ewBAI_2010 |
| lwBAI | 2011 | 2010 (one year) | 2012 (one year) | Rs_lwBAI_2010; Rc_lwBAI_2010; Rl_lwBAI_2010 |
| ewWD | 2010 | 2009 (one year) | 2011 (one year) | Rs_ewWD_2010; Rc_ewWD_2010; Rl_ewWD_2010 |
| lwWD | 2010 | 2009 (one year) | 2011 (one year) | Rs_lwWD_2010; Rc_lwWD_2010; Rl_lwWD_2010 |
| LwP | 2011 | 2010 (one year) | 2012 (one year) | Rs_LwP_2010; Rc_LwP_2010; Rl_LwP_2010 |
| ewBAI | 2013 | 2011-2012 (two years) | 2014-2015 (two years) | Rs_ewBAI_2012; Rc_ewBAI_2012; Rl_ewBAI_2012 |
| lwBAI | 2013 | 2012 (one year) | 2014 (one year) | Rs_lwBAI_2012; Rc_lwBAI_2012; Rl_lwBAI_2012 |
| ewWD | 2012 | 2011 (one year) | 2013 (one year) | Rs_ewWD_2012; Rc_ewWD_2012; Rl_ewWD_2012 |
| lwWD | 2012 | 2011 (one year) | 2013 (one year) | Rs_lwWD_2012; Rc_lwWD_2012; Rl_lwWD_2012 |
| LwP | 2013-2014 | 2012 (one year) | 2015 (one year) | Rs_LwP_2012; Rc_LwP_2012; Rl_LwP_2012 |
| ewBAI | 2016 | 2014-2015 (two years) | 2017-2018 (two years) | Rs_ewBAI_2015; Rc_ewBAI_2015; Rl_ewBAI_2015 |
| lwBAI | 2015 | 2014 (one year) | 2016 (one year) | Rs_lwBAI_2015; Rc_lwBAI_2015; Rl_lwBAI_2015 |
| ewWD | 2015-2016 | 2013-2014 (two years) | 2017-2018 (two years) | Rs_ewWD_2015; Rc_ewWD_2015; Rl_ewWD_2015 |
| lwWD | 2015-2016 | 2013-2014 (two years) | 2017-2018 (two years) | Rs_lwWD_2015; Rc_lwWD_2015; Rl_lwWD_2015 |
| LwP | 2016 | 2015 (one year) | 2017 (one year) | Rs_LwP_2015; Rc_LwP_2015; Rl_LwP_2015 |

**Table S5. Statistics obtained for time-series repeated-measures models fitted for radial growth and wood density.**

Statistics for complete time series for radial growth and wood density (BAI, WD), as well as earlywood (ewBAI, ewWD) and latewood (lwBAI, lwWD) series, are presented. In these time-series repeated measures models, the main effects of group and period, the nested family effect within group (fam:group), and interaction effects between period and group, as well as between the nested family effect within group and period, were treated as fixed effects. The random effects included the main effect of block, as well as interactions between period and year (Period:year), between block and year (year:block), between group and year (group:year), and between the nested family effect within group and year (year:fam(group)). Likelihood ratio tests were conducted using ASReml-R (lrt.asreml function) to assess the significance of each random effect by comparing the full model with a reduced model that excluded the random effect being tested. P-values of the likelihood ratio test are reported for random effects, while WALD statistics and associated p-values with their corresponding levels of significance are reported for fixed effects. *$P < 0.05$, **$P < 0.01$, ***$P < 0.001$

| | | **BAI** | | **ewBAI** | | **lwBAI** | | **WD** | | **ewWD** | | **lwWD** | | **LwP** | |
|---|---|---|---|---|---|---|---|---|---|---|---|---|---|---|---|
| Transformation type | | square root | | square root | | square root | | reciprocal | | reciprocal | | - | | square root | |
| Effect | Type | Wald | p-value | Wald | p-value | Wald | p-value | Wald | p-value | Wald | p-value | Wald | p-value | Wald | p-value |
| Intercept | - | 979.03 | <0.001*** | 271.00 | <0.001*** | 643.23 | <0.001*** | 4975.15 | <0.001*** | 3652.04 | <0.001*** | 18181.57 | <0.001*** | 1361.73 | <0.001*** |
| Group | Fixed | 3.09 | 0.38 | 120.12 | <0.001*** | 243.59 | <0.001*** | 68.02 | <0.001*** | 80.77 | <0.001*** | 20.24 | <0.001*** | 66.95 | <0.001*** |
| Period | Fixed | 0.26 | 0.61 | 1.97 | 0.16 | 11.07 | <0.001*** | 13.95 | <0.001*** | 23.20 | <0.001*** | 9.60 | 0.0019** | 6.15 | 0.013* |
| Period:group | Fixed | 16.87 | <0.001*** | 127.62 | <0.001*** | 44.33 | <0.001*** | 43.57 | <0.001*** | 5.82 | 0.12 | 4.92 | 0.18 | 66.85 | <0.001*** |
| Fam(group) | Fixed | 21.75 | 0.24 | 27.23 | <0.001*** | 62.62 | <0.001*** | 77.78 | <0.001*** | 116.05 | <0.001*** | 62.91 | <0.001*** | 74.54 | <0.001*** |
| Period:fam(group) | Fixed | 20.06 | 0.33 | 46.87 | <0.001*** | 22.99 | <0.001*** | 38.53 | 0.0033** | 28.61 | 0.053 | 11.22 | 0.88 | 33.34 | 0.015* |
| Period:year | Random | | <0.001*** | | <0.001*** | | <0.001*** | | <0.001*** | | <0.001*** | | <0.001*** | | <0.001*** |
| Block | Random | | <0.001*** | | <0.001*** | | <0.001*** | | <0.001*** | | <0.001*** | | <0.001*** | | <0.001*** |
| Year:block | Random | | <0.001*** | | <0.001*** | | 0.013* | | <0.001*** | | <0.001*** | | <0.001*** | | 0.0053** |
| Group:year | Random | | 0.5 | | <0.001*** | | <0.001*** | | <0.001*** | | <0.001*** | | <0.001*** | | <0.001*** |
| Year:fam(group) | Random | | 0.015* | | <0.001*** | | 0.062 | | <0.001*** | | <0.001*** | | <0.001*** | | <0.001*** |

**Table S6. Summary of taxonomic and family effects on variation in dendrometric traits.**

**(a)** In these models, group and block effects were treated as fixed effects, while family and the interaction between family and block effects were treated as random effects. P-values of the likelihood ratio test are reported for random effects, while WALD statistics and associated p-values with their corresponding levels of significance are reported for fixed effects. *$P < 0.05$, **$P < 0.01$, ***$P < 0.001$**.**
**(b)** Averages and differences between the groups for dendrometric traits. In this table, *Pma x Pru* and *Pru x Pma* represent the two hybrid groups, with *Pma* denoting pure black spruce trees and *Pru* denoting pure red spruce trees. Post-hoc statistical grouping of groups was conducted using Tukey's pairwise comparison method with a significance level of 0.05, and significant differences between groups are indicated by different letters.

**(a) Dendrometric traits model summaries**

| | | **H_10** | | **DBH_10** | | **H_20** | | **DBH_20** | |
|---|---|---|---|---|---|---|---|---|---|
| Effect | Type | Wald | p-value | Wald | p-value | Wald | p-value | Wald | p-value |
| Intercept | | 2223.23 | <0.001*** | 745.72 | <0.001*** | 3825.45 | <0.001*** | 2269.47 | <0.001*** |
| Group | Fixed | 30.06 | <0.001*** | 25.76 | <0.001*** | 14.60 | 0.0022** | 9.19 | 0.027* |
| Block | Fixed | 186.53 | <0.001*** | 345.68 | <0.001*** | 165.28 | <0.001*** | 85.50 | <0.001*** |
| Fam(group) | Random | | <0.001*** | | <0.001*** | | <0.001*** | | <0.001*** |
| Family:block | Random | | <0.001*** | | <0.001*** | | <0.001*** | | 0.019* |

**(b) Dendrometric traits group estimates, post-hoc comparisons and family effects**

| **Trait type** | **Trait name** | **Age** | ***Pma* Mean (SE)** | ***Pma x Pru* Mean (SE)** | ***Pru x Pma* Mean (SE)** | ***Pru* Mean (SE)** | **Group p-value** | **Family p-value** | **Family effect (%)** |
|---|---|---|---|---|---|---|---|---|---|
| Height | H_10 | 10 | 395 (15.0) a | 397 (13.9) a | 386 (18.8) a | 288 (16.5) b | <0.001*** | <0.001*** | 14.6 |
| Height | H_20 | 20 | 794 (24.1) a | 820 (22.3) a | 796 (30.0) ab | 683 (26.6) b | 0.0022** | <0.001*** | 25.4 |
| DBH | DBH_10 | 10 | 51.3 (3.31) a | 53.2 (3.06) a | 50.0 (4.13) a | 30.0 (3.63) b | <0.001*** | <0.001*** | 16.6 |
| DBH | DBH_20 | 20 | 123 (5.18) a | 138 (4.79) a | 135 (6.45) a | 116 (5.71) a | 0.027* | <0.001*** | 21.8 |

**Table S7. Results from linear models for climate-sensitivity traits.**

Detailed descriptions of each CS trait can be found in line 2. In these models, group and block effects were treated as fixed effects, while family and the interaction between family and block effects were considered random effects. Likelihood ratio tests were conducted using ASReml-R (lrt.asreml function) to assess the significance of each random effect by comparing the full model with a reduced model that excluded the random effect being tested. P-values of the likelihood ratio test are reported for random effects, while WALD statistics and associated p-values with their corresponding levels of significance are reported for fixed effects. **P* <0.05, ***P* <0.01, ****P* <0.001.

### (a) Model summaries part 1 of 3

| **Trait name** | | **ewBAI_SMI_PrevAut** | | **ewBAI_TMax_PrevAut** | | **ewBAI_LFR_Prev** | | **ewBAI_EFD** | | **ewBAI_TMin_Win** | |
|---|---|---|---|---|---|---|---|---|---|---|---|
| **Trait description** | | **Sensitivity of earlywood BAI to the soil moisture of the preceding autumn** | | **Sensitivity of earlywood BAI to the maximum temperature of the preceding autumn** | | **Sensitivity of earlywood BAI to the late frost risk of the preceding year** | | **Sensitivity of earlywood BAI to extreme frost days in the current year** | | **Sensitivity of earlywood BAI to minimal temperature in the current year** | |
| Effect | Type | Wald | p-value | Wald | p-value | Wald | p-value | Wald | p-value | Wald | p-value |
| Intercept | | 96.07 | <0.001*** | 156.35 | <0.001*** | 352.35 | <0.001*** | 110.24 | <0.001*** | 548.39 | <0.001*** |
| Group | Fixed | 65.40 | <0.001*** | 15.60 | 0.0014** | 11.90 | 0.0077** | 6.46 | 0.091 | 16.75 | <0.001*** |
| Block | Fixed | 19.55 | 0.67 | 25.89 | 0.31 | 26.98 | 0.26 | 37.96 | 0.026* | 42.45 | 0.008** |
| Fam(group) | Random | | 0.0072** | | <0.001** | | 0.0097** | | <0.001*** | | 0.0075** |
| Fam:Block | Random | | 0.071 | | 0.031* | | <0.001*** | | 0.033* | | 0.4 |

### (b) Model summaries part 2 of 3

| **Trait name** | | **ewWD_SMI_Sum** | | **ewWD_TMax_Sum** | | **ewWD_TMin_Win** | | **ewWD _WTD_Prev** | | **ewWD_SMI_PrevSpr** | | **ewWD_TMax_PrevSum** | |
|---|---|---|---|---|---|---|---|---|---|---|---|---|---|
| **Trait description** | | **Sensitivity of earlywood WD to the soil moisture of the current summer** | | **Sensitivity of earlywood WD to the maximal temperature of the current summer** | | **Sensitivity of earlywood WD to the minimal temperature of the current summer** | | **Sensitivity of earlywood WD to winter thaw days of the preceding year** | | **Sensitivity of earlywood WD to soil moisture of the preceding spring** | | **Sensitivity of earlywood WD to maximal temperature of the preceding summer** | |
| Effect | Type | Wald | p-value | Wald | p-value | Wald | p-value | Wald | p-value | Wald | p-value | Wald | p-value |
| Intercept | | 741.08 | <0.001*** | 584.58 | <0.001*** | 1434.56 | <0.001*** | 618.62 | <0.001*** | 192.47 | <0.001*** | 407.71 | <0.001*** |
| Group | Fixed | 4.88 | 0.18 | 8.45 | 0.038* | 81.45 | <0.001*** | 1.40 | 0.7 | 6.20 | 0.1 | 12.85 | 0.005** |
| Block | Fixed | 32.12 | 0.098 | 37.07 | 0.032* | 35.43 | 0.047* | 35.42 | 0.047* | 39.78 | 0.016* | 35.78 | 0.043* |
| Fam(group) | Random | | 0.5 | | 0.5 | | 0.42 | | 0.063 | | <0.001*** | | 0.027* |
| Fam:Block | Random | | 0.057 | | 0.017* | | 0.12 | | 0.5 | | 0.041* | | 0.085 |

### (c) Model summaries part 3 of 3

| Trait name | | lwWD_SMI_Sum | | lwWD _SMI_PrevSpr | | lwWD_WTD_Prev | | lwWD_TMax_PrevSum | | lwBAI_EFD | | LwP_TMax_PrevAut | |
|---|---|---|---|---|---|---|---|---|---|---|---|---|---|
| Trait description | | Sensitivity of latewood WD to soil moisture of the current summer | | Sensitivity of latewood WD to soil moisture of the preceding spring | | Sensitivity of latewood WD to winter thaw days of the preceding year | | Sensitivity of latewood WD to maximal temperature of the preceding summer | | Sensitivity of latewood BAI to extreme frost days in the current year | | Sensitivity of latewood percentage to maximal temperature of the preceding autumn | |
| Effect | Type | Wald | p-value | Wald | p-value | Wald | p-value | Wald | p-value | Wald | p-value | Wald | p-value |
| Intercept | | 699.07 | <0.001*** | 273.09 | <0.001*** | 291.70 | <0.001*** | 375.01 | <0.001*** | 21.50 | <0.001*** | 232.46 | <0.001*** |
| Group | Fixed | 10.88 | 0.012* | 8.51 | 0.037* | 38.13 | <0.001*** | 1.75 | 0.63 | 6.67 | 0.083 | 1.36 | 0.71 |
| Block | Fixed | 56.70 | <0.001*** | 45.69 | 0.0033** | 49.05 | 0.0012** | 55.41 | <0.001*** | 28.91 | 0.18 | 36.77 | 0.034* |
| Fam(group) | Random | | 0.27 | | 0.0018** | | 0.084 | | 0.18 | | 0.26 | | 0.5 |
| Fam:Block | Random | | 0.09 | | 0.44 | | 0.37 | | 0.17 | | 0.5 | | 0.066 |

**Table S8. Results obtained from linear models applied to resilience indices.**

Resilience components, including resistance (Rs), recovery (Rc), and resilience (Rl) of earlywood and latewood series for radial growth (ewBAI and lwBAI, (a) and (b) respectively) and wood density (ewWD and lwWD, (c) and (d) respectively), and latewood percentage (LwP, (e)). The resilience indices were calculated for both drought episodes (denoted by the extensions 2010 or 2012 at the end of the resilience component name) and the cold episode (denoted by the extension 2015 at the end of the name). Group and block effects were treated as fixed effects in the fitted models, while family and the interaction between family and block effect were considered random effects. Likelihood ratio tests were conducted using ASReml-R (lrt.asreml function) to assess the significance of each random effect by comparing the full model with a reduced model that excluded the random effect being tested. P-values of the likelihood ratio test are reported for random effects, while WALD statistics and associated p-values with their corresponding levels of significance are reported for fixed effects (*P <0.05, **P <0.01, ***P <0.001).

| **(a)** | | Rs_ewBAI_2010 | | Rc_ewBAI_2010 | | Rl_ewBAI_2010 | | Rs_ewBAI_2012 | | Rc_ewBAI_2012 | | Rl_ewBAI_2012 | | Rs_ewBAI_2015 | | Rc_ewBAI_2015 | | Rl_ewBAI_2015 | |
|---|---|---|---|---|---|---|---|---|---|---|---|---|---|---|---|---|---|---|---|
| Effect | Type | Wald | p-value | Wald | p-value | Wald | p-value | Wald | p-value | Wald | p-value | Wald | p-value | Wald | p-value | Wald | p-value | Wald | p-value |
| Intercept | | 6543.48 | <0.001*** | 2716.10 | <0.001*** | 7418.50 | <0.001*** | 8734.86 | <0.001*** | 11043.82 | <0.001*** | 13409.02 | <0.001*** | 7370.10 | <0.001*** | 5527.18 | <0.001*** | 12047.59 | <0.001*** |
| Group | Fixed | 21.29 | <0.001*** | 10.73 | 0.013* | 4.07 | 0.25 | 14.95 | 0.0019** | 53.78 | <0.001*** | 13.16 | 0.0043** | 6.31 | 0.097 | 19.05 | <0.001*** | 60.58 | <0.001*** |
| Block | Fixed | 35.20 | 0.05* | 38.55 | 0.022* | 36.15 | 0.04* | 49.87 | <0.001*** | 32.92 | 0.083 | 43.95 | 0.0053** | 62.90 | <0.001*** | 33.76 | 0.069 | 36.48 | 0.037* |
| Fam(group) | Random | | 0.0057** | | <0.001*** | | 0.017* | | <0.001*** | | 0.017* | | 0.0098** | | 0.019* | | 0.0036** | | 0.054 |
| Fam:Block | Random | | 0.0046** | | 0.023* | | 0.0014** | | 0.002** | | 0.017* | | 0.16 | | <0.001*** | | 0.12 | | 0.12 |

| **(b)** | | Rs_lwBAI_2010 | | Rc_lwBAI_2010 | | Rl_lwBAI_2010 | | Rs_lwBAI_2012 | | Rc_lwBAI_2012 | | Rl_lwBAI_2012 | | Rs_lwBAI_2015 | | Rc_lwBAI_2015 | | Rl_lwBAI_2015 | |
|---|---|---|---|---|---|---|---|---|---|---|---|---|---|---|---|---|---|---|---|
| Effect | Type | Wald | p-value | Wald | p-value | Wald | p-value | Wald | p-value | Wald | p-value | Wald | p-value | Wald | p-value | Wald | p-value | Wald | p-value |
| Intercept | | 4.91 | 0.027* | 21.53 | <0.001*** | 2.33 | 0.13 | 8.44 | 0.0037** | 0.01 | 0.92 | 11.14 | <0.001*** | 11.57 | <0.001*** | 34.60 | <0.001*** | 8.59 | 0.0034** |
| Group | Fixed | 0.90 | 0.82 | 14.29 | 0.0025** | 6.20 | 0.1 | 12.82 | 0.005** | 4.55 | 0.21 | 7.77 | 0.051 | 2.38 | 0.5 | 3.77 | 0.29 | 4.33 | 0.23 |
| Block | Fixed | 8.75 | 1.00 | 32.30 | 0.094 | 21.63 | 0.54 | 28.53 | 0.2 | 30.68 | 0.13 | 47.10 | 0.0022** | 32.06 | 0.099 | 24.22 | 0.39 | 23.47 | 0.43 |
| Fam(group) | Random | | 0.41 | | 0.5 | | 0.31 | | 0.5 | | 0.094 | | 0.43 | | 0.3 | | 0.32 | | 0.5 |
| Fam:Block | Random | | <0.001*** | | 0.5 | | 0.0035** | | 0.0013** | | 0.026* | | 0.0055** | | 0.058 | | 0.023* | | 0.0043** |

| (c) | | Rs_ewWD_2010 | | Rc_ewWD_2010 | | Rl_ewWD_2010 | | Rs_ewWD_2012 | | Rc_ewWD_2012 | | Rl_ewWD_2012 | | Rs_ewWD_2015 | | Rc_ewWD_2015 | | Rl_ewWD_2015 | |
|---|---|---|---|---|---|---|---|---|---|---|---|---|---|---|---|---|---|---|---|
| Effect | Type | Wald | p-value | Wald | p-value | Wald | p-value | Wald | p-value | Wald | p-value | Wald | p-value | Wald | p-value | Wald | p-value | Wald | p-value |
| Intercept | | 119738.30 | <0.001*** | 73170.07 | <0.001*** | 108573.01 | <0.001*** | 63985.24 | <0.001*** | 170237.66 | <0.001*** | 167269.25 | <0.001*** | 101965.41 | <0.001*** | 167696.13 | <0.001*** | 372261.88 | <0.001*** |
| Group | Fixed | 35.71 | <0.001*** | 1.91 | 0.59 | 16.39 | <0.001*** | 8.79 | 0.032* | 40.38 | <0.001*** | 3.37 | 0.34 | 41.53 | <0.001*** | 43.87 | <0.001*** | 5.04 | 0.17 |
| Block | Fixed | 38.63 | 0.022* | 24.51 | 0.38 | 55.83 | <0.001*** | 43.26 | 0.0065** | 18.48 | 0.73 | 55.66 | <0.001*** | 43.24 | 0.0065** | 37.15 | 0.031* | 18.86 | 0.71 |
| Fam(group) | Random | | 0.019* | | <0.001*** | | 0.0066** | | <0.001*** | | 0.11 | | 0.047* | | <0.001*** | | 0.0062** | | 0.035* |
| Fam:Block | Random | | 0.5 | | 0.21 | | 0.058 | | 0.31 | | 0.5 | | 0.36 | | 0.016* | | 0.22 | | 0.074 |

| (d) | | Rs_lwWD_2010 | | Rc_lwWD_2010 | | Rl_lwWD_2010 | | Rs_lwWD_2012 | | Rc_lwWD_2012 | | Rl_lwWD_2012 | | Rs_lwWD_2015 | | Rc_lwWD_2015 | | Rl_lwWD_2015 | |
|---|---|---|---|---|---|---|---|---|---|---|---|---|---|---|---|---|---|---|---|
| Effect | Type | Wald | p-value | Wald | p-value | Wald | p-value | Wald | p-value | Wald | p-value | Wald | p-value | Wald | p-value | Wald | p-value | Wald | p-value |
| Intercept | | 80826.40 | <0.001*** | 106525.05 | <0.001*** | 59564.69 | <0.001*** | 70196.04 | <0.001*** | 99810.27 | <0.001*** | 127469.93 | <0.001*** | 74495.77 | <0.001*** | 93344.08 | <0.001*** | 382600.44 | <0.001*** |
| Group | Fixed | 34.53 | <0.001*** | 4.03 | 0.26 | 14.80 | 0.002** | 4.22 | 0.24 | 23.43 | <0.001*** | 14.68 | 0.0021** | 17.59 | <0.001*** | 18.89 | <0.001*** | 10.44 | 0.015* |
| Block | Fixed | 20.72 | 0.6 | 27.13 | 0.25 | 25.53 | 0.32 | 41.49 | 0.01* | 24.84 | 0.36 | 25.46 | 0.33 | 28.43 | 0.2 | 35.51 | 0.046* | 26.53 | 0.28 |
| Fam(group) | Random | | 0.29 | | 0.5 | | 0.021* | | 0.056 | | 0.26 | | 0.5 | | 0.0043** | | 0.019* | | 0.5 |
| Fam:Block | Random | | 0.46 | | 0.35 | | 0.5 | | 0.5 | | 0.49 | | 0.5 | | 0.5 | | 0.5 | | 0.087 |

| (e) | | Rs_LwP_2010 | | Rc_LwP_2010 | | Rl_LwP_2010 | | Rs_LwP_2012 | | Rc_LwP_2012 | | Rl_LwP_2012 | | Rs_LwP_2015 | | Rc_LwP_2015 | | Rl_LwP_2015 | |
|---|---|---|---|---|---|---|---|---|---|---|---|---|---|---|---|---|---|---|---|
| Effect | Type | Wald | p-value | Wald | p-value | Wald | p-value | Wald | p-value | Wald | p-value | Wald | p-value | Wald | p-value | Wald | p-value | Wald | p-value |
| Intercept | | 22.38 | <0.001*** | 28.28 | <0.001*** | 0.97 | 0.32 | 5.14 | 0.023* | 74.12 | <0.001*** | 42.31 | <0.001*** | 96.22 | <0.001*** | 7.21 | 0.0073** | 123.93 | <0.001*** |
| Group | Fixed | 2.91 | 0.41 | 17.71 | <0.001*** | 7.50 | 0.058 | 16.88 | <0.001*** | 24.56 | <0.001*** | 2.60 | 0.46 | 9.27 | 0.026* | 5.26 | 0.15 | 10.07 | 0.018* |
| Block | Fixed | 9.33 | 0.99 | 29.04 | 0.18 | 26.39 | 0.28 | 30.16 | 0.14 | 40.31 | 0.014* | 30.15 | 0.15 | 24.43 | 0.38 | 28.04 | 0.21 | 36.60 | 0.036* |
| Fam(Group) | Random | | 0.42 | | 0.061 | | 0.26 | | 0.13 | | 0.5 | 42.31 | 0.5 | | 0.092 | | 0.5 | | 0.5 |
| Fam:Block | Random | | <0.001*** | | 0.12 | | 0.0042** | | <0.001*** | | 0.018* | 2.60 | 0.037* | | 0.12 | | 0.12 | | 0.5 |

**Table S9. Results of linear modeling for resilience, provided for the three identified climatic stress periods during the study period.**

Resilience (Rl) was calculated from detrended data of traits from both earlywood and latewood series of radial growth and wood density, in the same fashion as the two other components presented in Table 4. Significance levels for group and family effects are indicated as follows: *$P < 0.05$, **$P < 0.01$, and ***$P < 0.001$. In cases where the group effect was significant ($P < 0.05$), Tukey's pairwise comparison method was used for post-hoc grouping of groups with a significance level of 0.05, and significant differences between groups are indicated by different letters. If needed, logarithmic transformations were applied to respect data normality. Trait definitions: ewBAI, earlywood basal area increment; lwBAI, latewood basal area increment; ewWD, earlywood wood density; lwWD, latewood wood density; LwP, latewood percentage.

| Associated climatic stress period | Resilience trait | Family effect | Group effect | *Pma*; Mean (SE) | *Pma x Pru*; Mean (SE) | *Pru x Pma*; Mean (SE) | *Pru*; Mean (SE) |
|---|---|---|---|---|---|---|---|
| 2010 summer drought | ewBAI | 0.017* | 0.25 | 1.11 (0.024) | 1.05 (0.022) | 1.07 (0.0308) | 1.13 (0.0276) |
| | ewWD | 0.0066** | <0.001*** | 1.03 (0.00573)a | 1.01 (0.00526)a | 1.02 (0.00758)a | 0.985 (0.00709)b |
| | lwBAI | 0.31 | 0.1 | 0.945 (0.0458) | 1.05 (0.0462) | 1.07 (0.0653) | 1.14 (0.0648) |
| | lwWD | 0.021* | 0.002** | 0.979 (0.00757)b | 0.987 (0.00694)b | 0.985 (0.00995)ab | 1.02 (0.00953)a |
| | LwP | 0.26 | 0.058 | 0.876 (0.0439) | 0.997 (0.0454) | 1.01 (0.0639) | 1.07 (0.0628) |
| 2012 summer drought | ewBAI | 0.0098** | 0.0043** | 1.02 (0.0162)a | 0.965 (0.0149)ab | 0.936 (0.0203)b | 1.00 (0.0183)ab |
| | ewWD | 0.047* | 0.34 | 0.991 (0.00468) | 1.00 (0.00428) | 0.998 (0.00587) | 1.00 (0.00532) |
| | lwBAI | 0.43 | 0.051 | 1.02 (0.0416) | 0.942 (0.0356) | 0.855 (0.043) | 0.867 (0.0416) |
| | lwWD | 0.5 | 0.0021** | 0.993 (0.00538)a | 0.989 (0.00485)a | 0.997 (0.00653)a | 0.965 (0.00624)b |
| | LwP | 0.5 | 0.46 | 0.911 (0.0336) | 0.853 (0.029) | 0.858 (0.0385) | 0.885 (0.0383) |
| Cold conditions from 2014 to 2016 | ewBAI | 0.054 | <0.001*** | 0.828 (0.0161)c | 0.930 (0.0149)b | 0.919 (0.02)b | 1.01 (0.0181)a |
| | ewWD | 0.035* | 0.17 | 1.01 (0.00316) | 1.01 (0.00294) | 1.01 (0.00394) | 1.00 (0.00356) |
| | lwBAI | 0.5 | 0.23 | 1.10 (0.0441) | 1.01 (0.0378) | 1.13 (0.0558) | 1.03 (0.049) |
| | lwWD | 0.5 | 0.015* | 1.03 (0.00327)b | 1.04 (0.003)a | 1.04 (0.00391)a | 1.04 (0.00374)ab |
| | LwP | 0.5 | 0.018* | 1.29 (0.0463)a | 1.24 (0.0415)a | 1.30 (0.0565)a | 1.13 (0.0465)a |
